# An icosahedral virus as a fluorescent calibration standard: a method for counting protein molecules in cells by fluorescence microscopy

**DOI:** 10.1101/088617

**Authors:** John M. Murray

## Abstract

The ability to replace genes coding for cellular proteins with DNA that codes for fluorescent protein-tagged versions opens the way to counting the number of molecules of each protein component of macromolecular assemblies *in vivo* by measuring fluorescence microscopically. Converting fluorescence to absolute numbers of molecules requires a fluorescent standard whose molecular composition is known precisely. In this report the construction, properties, and mode of using a set of fluorescence calibration standards are described. The standards are based on an icosahedral virus engineered to contain exactly 240 copies of one of seven different fluorescent proteins. Two applications of the fluorescent standards to counting molecules in the human parasite *Toxoplasma gondii* are described. Methods for improving the preciseness of the measurements and minimizing potential inaccuracies are emphasized.

**Lay Abstract:** A broad goal of modern biology is to understand how the machines within living cells work. It is nowadays routine to identify the individual protein components of a machine, but not yet straightforward to tell how many copies of each component are needed to build a functional assembly. In many types of cells it is now possible to substitute for the native proteins within cells altered versions that are fluorescent. If one knew how much fluorescence is generated by a single molecule of the altered protein, then one could use a light microscope to count the number of copies of the protein in a cellular machine by simply measuring the total fluorescence coming from that part of the cell. This paper describes the construction and methods for using a set of fluorescent virus particles that can be used to determine how much fluorescence is contributed by one molecule of fluorescent protein. The virus particles were chosen for this role because the particular icosahedral symmetry of their structure guarantees that each particle contains exactly 240 copies of one fluorescent protein.

## Introduction

An extremely powerful tool for cell biologists became available with the advent of the fluorescent proteins (FPs) (Giepmans et al., 2006). With this tool, and with the ability to engineer in many organisms homologous replacements of endogenous genes with FP-tagged versions of those genes, it should in principle be possible to count the absolute number of molecules of a specific protein in a living cell, by measuring the amount of fluorescence emitted by the FP-tagged protein. Many examples of this application have been reported (for review, see (Verdaasdonk et al., 2014))

In order to convert the measured fluorescence into the absolute number of molecules, some sort of calibration standard is required. For the most part, the different attempts to count molecules in this way have each used a different means of calibration. It would be useful for cell biologists to have a set of shared calibration standards that could be used by anyone for comparing results from different labs. An ideal shared standard would have at least these properties: wide accessibility, convenience, reproducibility, and long-term stability. It should also have well characterized and documented properties, and be extensible to new fluorescent proteins as they become available. I report here the production and properties of a candidate set of fluorescent standards, based on an alphavirus particle whose genome has been modified to express an FP-tagged version of a capsid protein. The particles are bright, stable, and have ultimate reproducibility, as they are self-replicating entities. Two examples of the application of these standards to counting proteins in cells are described.

Accurate counting presupposes accurate measurement of fluorescence, and because the motivation for counting is frequently to compare two counts, the preciseness of counting is also of central concern. I report an extended analysis of the sources of and scale of inaccuracies and limits to preciseness that are inherent in using FPs in this way to count molecules in cells, and some methodological recommendations to minimize those errors.

## Results

### Construction of fluorescent Sindbis virus

Virus particles of known 3D structure have several advantages to recommend them as platforms for constructing fluorescence standards. The particles are small, stable, well-characterized, identical, self-replicating, and simple to prepare in a quantity that is very large compared to what is needed. Sindbis virus is a small (~60 nm) spherical alphavirus (Lloyd, 2009) that includes an 11.7 kb single-stranded RNA enclosed in an inner protein shell (nucleocapsid) made from the virally encoded CP protein, and an outer protein shell formed from 80 glycoprotein spikes, each spike containing three copies of a heterodimer assembled from viral proteins E1 and E2. A lipid bilayer derived from the host membrane lies between the two protein shells. The outer glycoprotein envelope has T=4 icosahedral symmetry; i.e., there are exactly 240 copies of the E1-E2 heterodimer per virion (Fuller, 1987). Several fluorescently tagged versions of Sindbis envelope glycoproteins have been described, and a 3D structure of mCherryFP-E2 Sindbis determined by cryoEM was recently published (Jose et al., 2015), demonstrating that each mature virus particle contains 240 copies of the mCherryFP-E2 fusion protein as expected.

Plasmids encoding Sindbis virus genomes with FP-tagged E2 were constructed for eight commonly used fluorescent proteins: mAppleFP, mCerulean3FP, mCherryFP, EGFP, mEmeraldFP, mNeonGreenFP, tdTomatoFP, and mVenusFP). The constructs differ from previously published FP-E2 fusion proteins only by the short flexible linkers used to couple the C-terminus of the FP to the N-terminus of E2 (AAAGSG), and C-terminus of E3 to N-terminus of the FP prior to furin cleavage (GAPGSA). Monolayer cultures of BHK-21 or Vero cells transfected with RNA produced by *in vitro* transcription from any of the constructed plasmid genomes (Rice et al., 1987) showed scattered brightly fluorescent single cells or small groups of cells within 6 hours after transfection (Figure 1). For seven of the eight constructs, the entire monolayer became fluorescent 24-36 hours after transfection, and infectious virus particles became abundant in the culture supernatant, with titers of 10^3^–10^7^/mL. Two of the constructs (EGFP and tdTomatoFP) readily produced fluorescent single cells after transfection with RNA, but rarely did the fluorescence spread to neighboring cells, and culture supernatants contained <10 pfu/mL of infectious virions.

**Figure 1.**
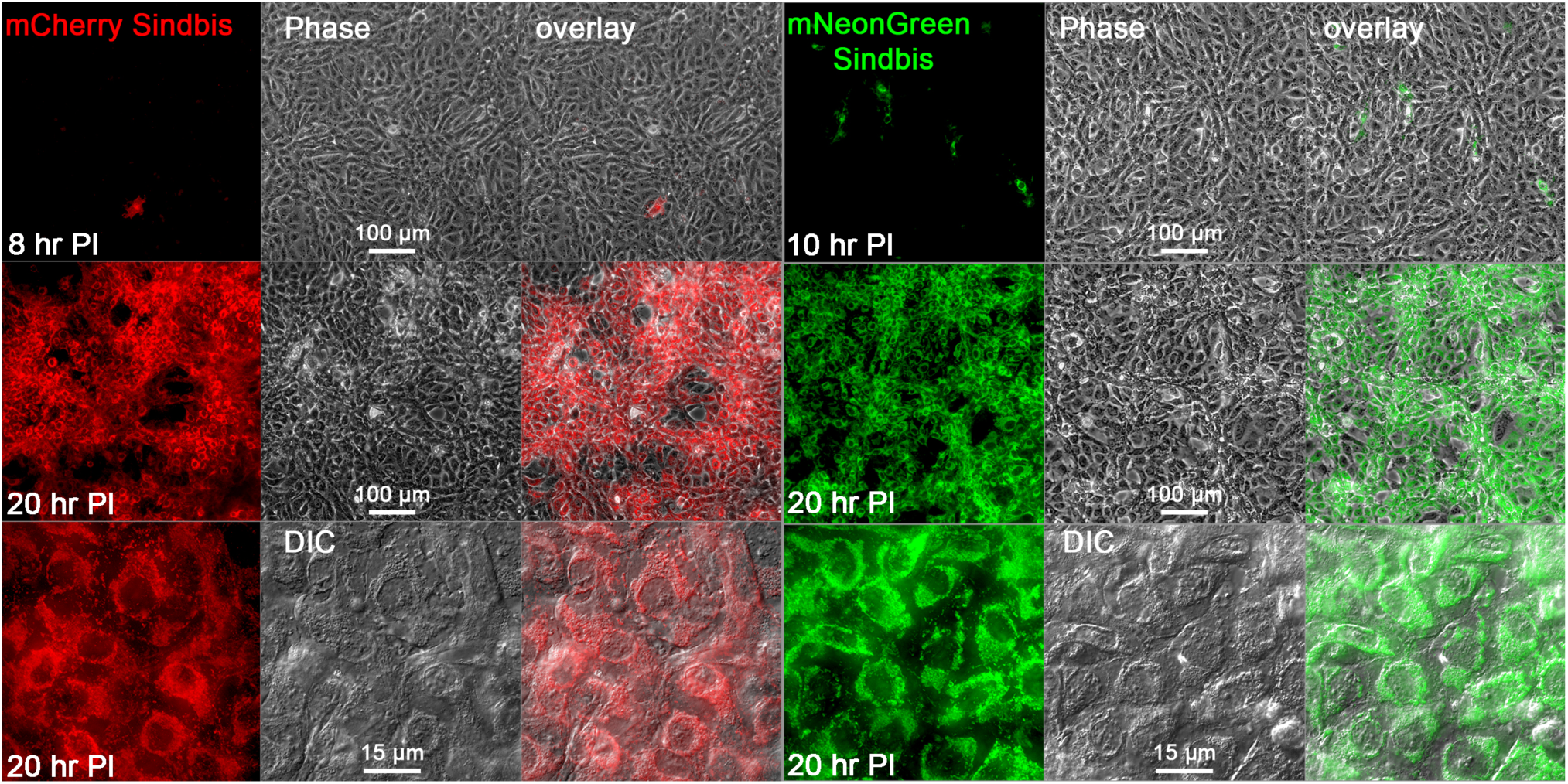
Infection of Vero cells with fluorescent Sindbis virus. (Left) Fluorescence and phase contrast or DIC images of a monolayer culture of cells 8 and 20 hours post-inoculation with mCherryFP Sindbis. The lower row shows higher magnification fluorescence and DIC images of the cells at 20 hours post-inoculation. (Right) Images of a monolayer culture of cells 10 and 20 hours post-inoculation with mNeonGreenFP Sindbis.

The failure to generate an infectious viral particle incorporating tdTomatoFP-E2 is perhaps not surprising, as tdTomatoFP is a tandem dimer, twice the size of the other FPs used here, and huge compared to the small E3 protein (476 versus 64 amino acid residues) it would replace in the maturing viral particles. However, given the success with the other six FPs, the failure with EGFP was both a surprise and puzzling. EGFP has been used in live cell imaging applications far more often than any other FP, and has become something of a gold standard against which the performance of other FPs in fusion proteins is assessed. At first glance, the effective FPs seemed to have little in common to distinguish them from EGFP. Some of the effective FPs were color variants differing in only a few amino acids from the *Aequorea victoria* derived EGFP such as mVenusFP and mCerulean3FP, but equally effective were *Discosoma sp* and *Branchiostoma lanceolatum* derived proteins such as mAppleFP, mCherryFP, and mNeonGreenFP, which have very little sequence homology to EGFP. One characteristic that does distinguish the effective FPs from EGFP is the presence in the former of dimerization-blocking mutations such as the A206K or L221K substitutions in the *Aequorea victoria* derived proteins. Although EGFP has only a weak tendency to dimerize and typically behaves well as a fusion partner in living cells, its residual dimerization capacity is higher than other more modern derivatives, as can be observed in an assay that directly measures tendency to oligomerize *in vivo* (Zacharias et al., 2002) (Costantini et al., 2012) (Cranfill et al., 2016). Swapping the EGFP coding sequence in EGFP-Sindbis with a mutant changed by a single amino acid residue, the monomerizing mutation L221K (Zacharias et al., 2002), produced a construct that yielded titers of infectious virions comparable to the other fluorescent Sindbis virions, confirming that even the rather weak dimerizing tendency of standard EGFP (L221) is sufficient to interfere with assembly of infectious virus particles.

### Characterization of fluorescent Sindbis virus

Centrifugation of culture media from cultures of cells infected with fluorescent Sindbis virus to remove cellular debris, and examination of the supernatant by fluorescence microscopy reveals numerous sub-resolution fluorescent dots (Figure 2). The virus suspensions are stable for many months when stored in buffer at 4°C. For longer term storage, the virus suspensions can be frozen and stored at -80°C for years with little loss. A few µL of frozen virus suspension scraped from the surface of a frozen aliquot is sufficient to infect a flask of host cells, yielding many mL of concentrated virus suspension in a few days. Serial re-amplification leads to takeover of the population by mutant dark forms after 5−10 passages, but as only a few µL of the original stock are needed for each amplification, and only 10−20 µL of the amplified product are needed for imaging, the supply of fluorescent virus, once made by transfection of host cells with *in vitro* transcribed viral RNA (see Methods), is in practice inexhaustible.

**Figure 2.**
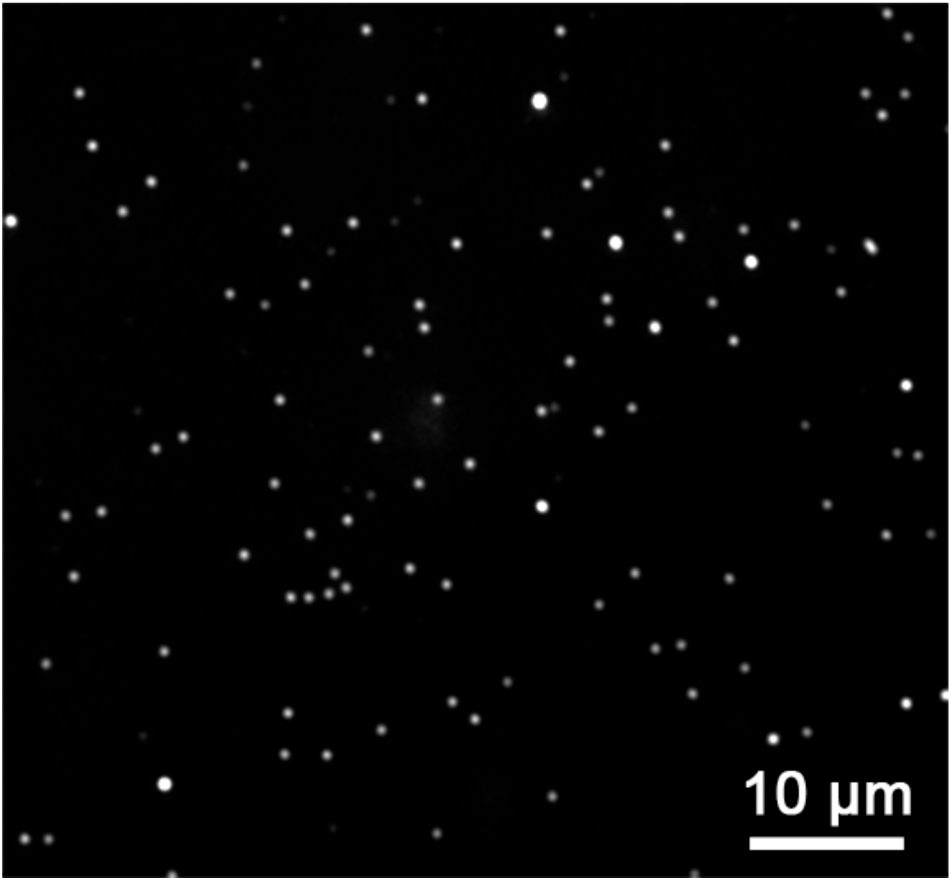
Epifluorescence microscope image of mCherryFP Sindbis virions

Quantitative measurements on a large number of fluorescent spots from cultures of different color variants show unimodal, or sometimes bimodal distributions with one large peak and a much smaller second peak of twice the brightness, presumably corresponding to single virions and unresolved pairs (Figure 3).

**Figure 3.**
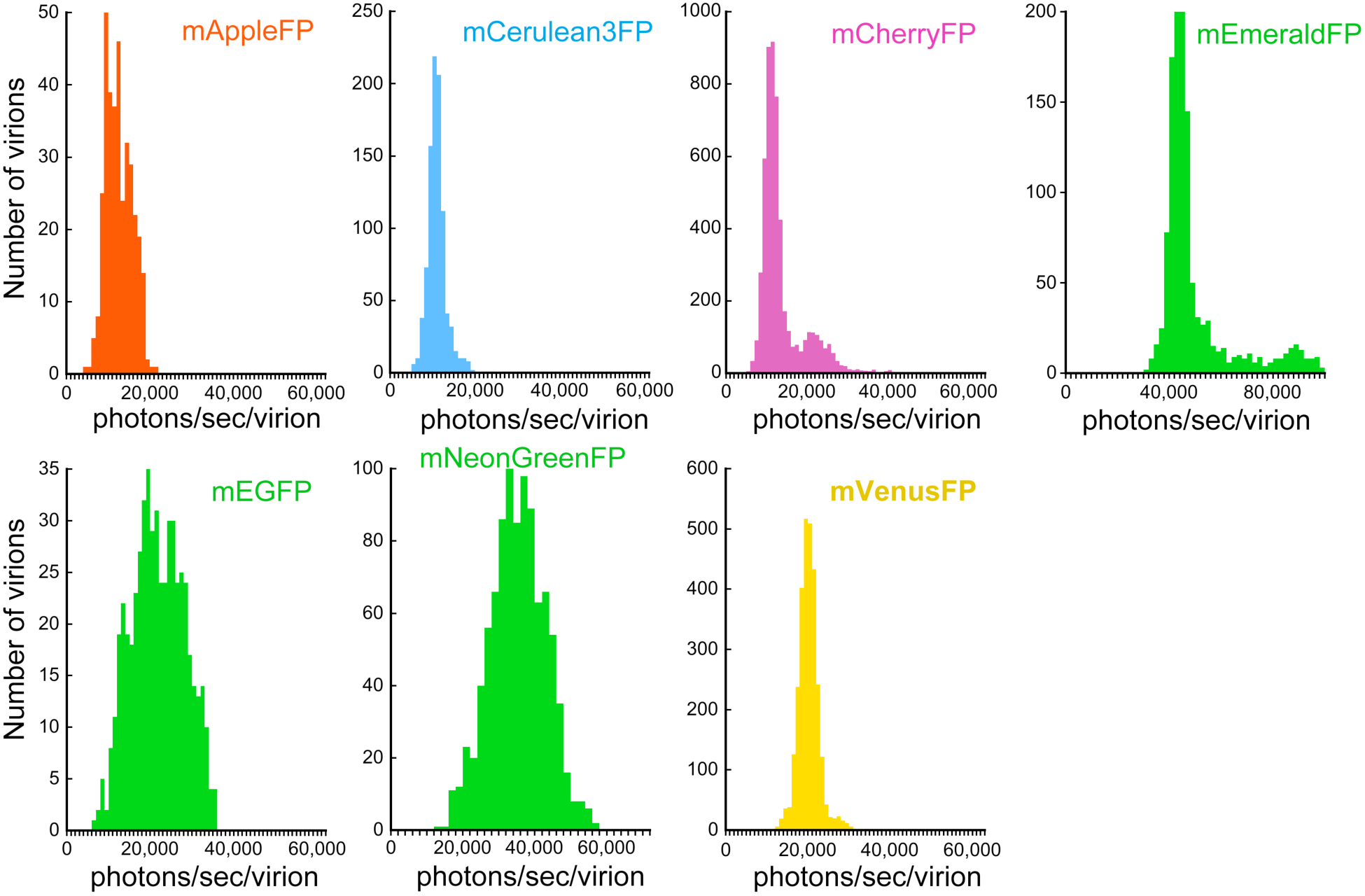
Histograms of measured brightness of various fluorescent Sindbis viruses. Brightness is given as detected photons per second per virus particle, net after background correction. The usual data curation (*see Methods*) was modified in order to reveal the small secondary peak of unresolved dimers in the mCherryFP-Sindbis and mEmeraldFP-Sindbis histograms.

### Analysis of the preciseness of measured brightness

In the images of fluorescent Sindbis virions shown in Figure 2, each virion has contributed 1−4 × 10^4^ photons. Ideally, the preciseness of measurement of the brightness of an individual virus particle would be limited only by Poisson noise, which in this case would be at most 1%. However, as the histograms in Figure 3 show, the variation in measured brightness among individual virus particles in a large population is 10−30 fold larger than this ideal for all of the color variants. This was a cause for concern, since it opened the possibility that there was some intrinsic heterogeneity in the population. If the number of fluorescent capsid protein molecules incorporated into each virion were not precisely 240 as expected on the basis of their icosahedral symmetry, but instead varied from virion to virion, then the accuracy of molecular counting utilizing these engineered viruses as fluorescent standards would be compromised. To address this possibility, an extensive set of experimental analyses and computational modeling was carried out.

Where is the “extra” noise coming from? One possibility is that the microscope/light source/camera system is introducing noise. To investigate this possibility, the amount of noise present in images of three radically different objects was determined. First, light from a battery powered (to avoid line-frequency fluctuations) incandescent source, diffused through a strongly scattering oil-in-water emulsion (diluted cream), was directed to the camera through the microscope optics. One hundred images were acquired in rapid succession and the pixel values in a small region were summed (Figure 4). The same measurements were carried out using light from the microscope mercury arc illumination system reflected off a mirror, and also using the fluorescence generated in a “lake” of fluorophore in aqueous solution. In all three cases, the variance in recorded photons was as predicted for simple Poisson noise. Thus any noise contribution from the imaging/detection system is insignificant compared to the inescapable Poisson noise in these images.

**Figure 4.**
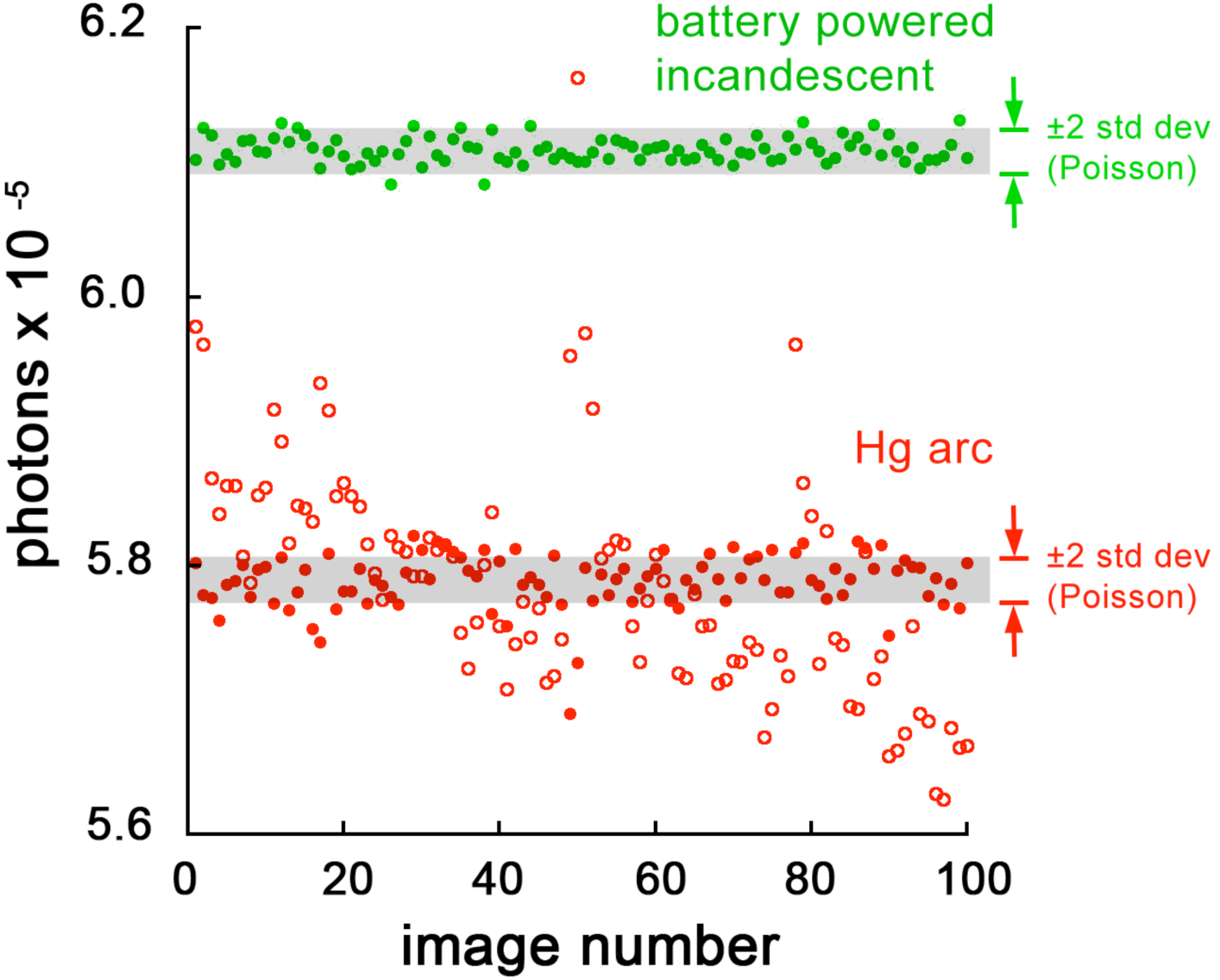
Imaging system noise level. The graph shows measurements of the summed pixel values in a 7 × 7 pixel box from 100 sequential images. The source of light was either a battery powered incandescent bulb (flashlight) viewed through a strongly scattering medium (*green*) or the output from the microscope’s Hg arc lamp reflected from a mirror in front of the objective lens (*red*). For the Hg arc, open circles show the raw measurements, closed circles show the measurements after correction for fluctuations in the Hg arc output, measured by a photosensor. The shaded bars show the expected range of variation due solely to Poisson noise.

One common source of brightness variation in fluorescence images is non-uniform illumination across the field of view. Figure 5 shows a typical example, an image of a thin layer of solution of a fluorophore, with intensities scaled relative to a value of 1.0 for the maximum. The three blue rings mark the contours at 90, 80, and 70% of maximum brightness. The center of symmetry of the illumination pattern, marked with an “X”, is offset from the center of the field of view of the camera (small red circle), which in turn is slightly offset from the optical axis (larger green circle) of the imaging system. The optic axis of the imaging system was defined empirically as the position in the field of view whose location in the image remains unshifted when the microscope’s accessory 1.5x lens is introduced or removed. This change of magnification causes a displacement in the image of all other points by an amount that is proportional to their distance from the optic axis.

**Figure 5.**
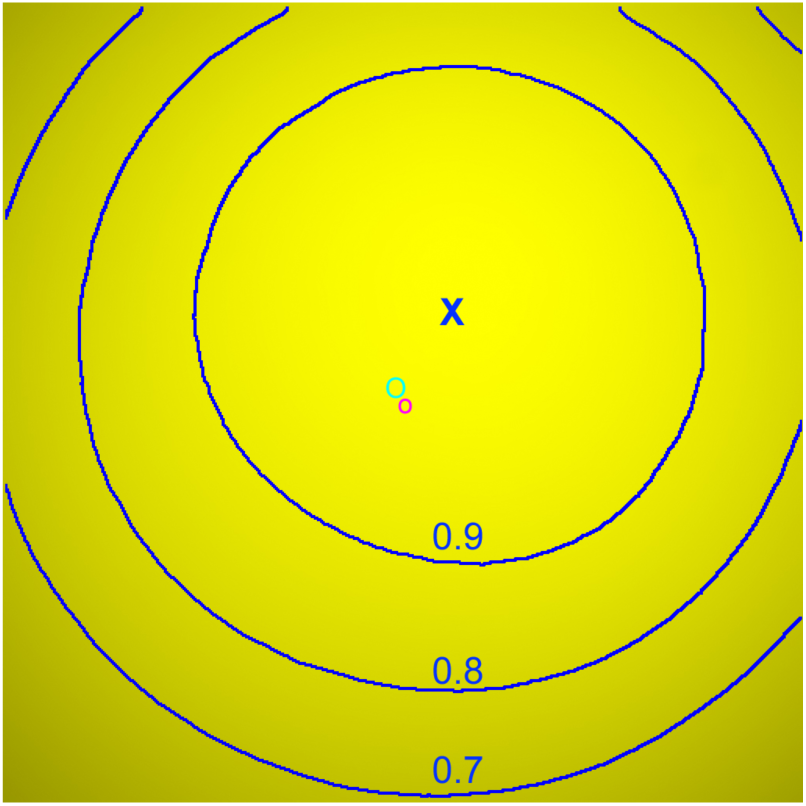
Illustration of typical non-uniform illumination. An image of a uniformly fluorescent sample was scaled relative to its maximum value, set to 1.0. Blue rings mark contours at 90, 80, and 70% of maximum brightness. “X”, center of symmetry of the illumination pattern; small red circle, center of the image; larger green circle, optical axis of the imaging system.

“Flat-fielding”, dividing every image by a scaled (maximum = 1.0) image of an object known to be uniformly fluorescent, is a common way of compensating for non-uniformity in illumination. The images of fluorescent Sindbis virions were flat-fielded in this way before analysis, and in addition the region of the images used for analysis was restricted to the area inside the 85% contour. The effectiveness of this procedure can be judged from Figure 6, a plot of measured brightness of fluorescent Sindbis virions before and after flat-fielding versus radial distance from the center of the illumination pattern. There is no significant correlation of measured brightness with radius after the flat-fielding correction. Thus the unexpectedly large variation in measured brightness among fluorescent virions is not due to ineffective compensation for non-uniform illumination. Although flat-fielding works reasonably well in this instance, a more robust solution to the problem is to assess brightness of the virions utilizing a quantitative measure that is relatively insensitive to illumination intensity as described below.

**Figure 6.**
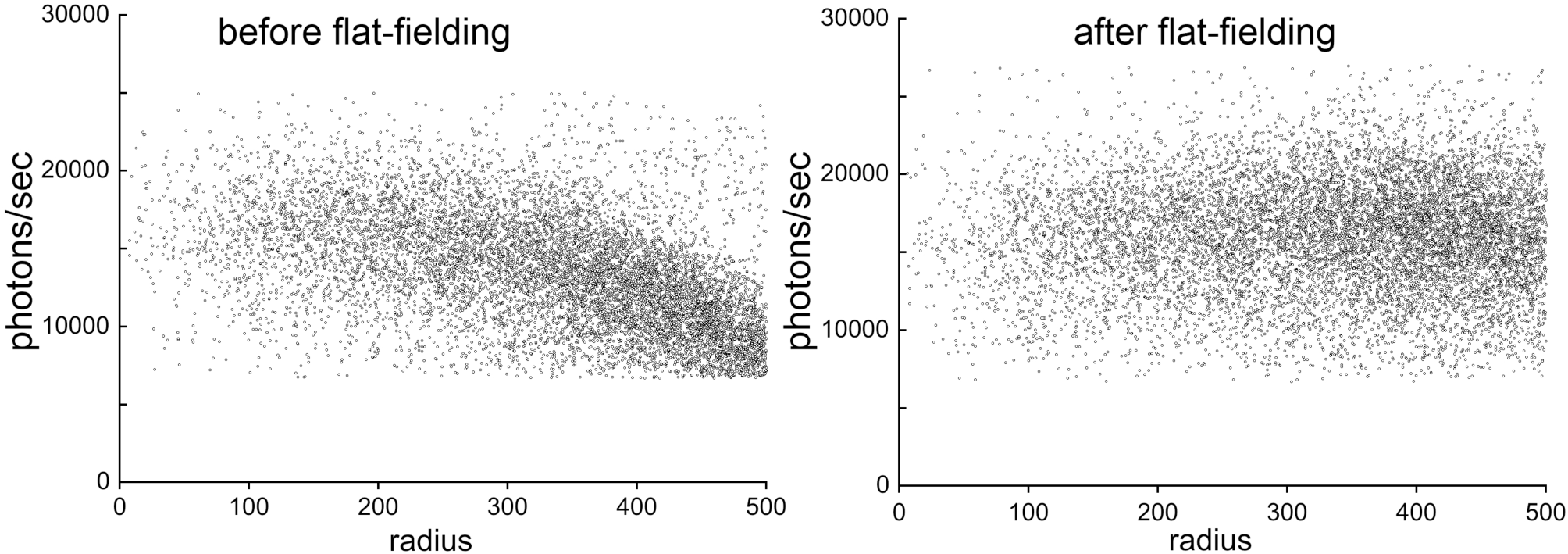
Plot of measured brightness of fluorescent virions *versus* radial distance from the center of symmetry of the illumination pattern. The pronounced fall-off of measured brightness with radius is removed by flat-fielding.

Several other possible causes of variable brightness were considered and rejected. (1) Many of the images of fluorescent virions were collected using an oil immersion lens with NA larger than the refractive index of the aqueous buffer in which the samples were mounted. Thus, in addition to the far-field illumination bathing the virions adsorbed to the coverslip, there will have been an evanescent field whose intensity decayed exponentially with distance from the coverslip-buffer interface. Conceivably, there might be small differences in the distance of each virion from the coverslip, which would be manifested as differences in intensity of the evanescent field, and hence differences in the amount of fluorescence generated from each virion. Evanescent field effects were ruled out by imaging with a water immersion lens, which cannot generate an evanescent field at the coverslip-buffer interface. The “extra” variance in the images acquired with the water immersion objective was not reduced compared to images acquired with an oil immersion objective. (2) The photophysical properties of the fluorescent proteins are complex (Day and Davidson, 2014; Shaner, 2014). Transient “blinking” of emitted fluorescence has been described, but, this is a stochastic property of single FP molecules, and therefore seemed unlikely to have much effect on the aggregated fluorescent output from an ensemble of 240 molecules unless there were some completely unknown mechanism that synchronized their blinking. However, other peculiarities such as reversible photobleaching, and rapid initial decay of brightness followed by much slower “normal” photobleaching, have also been described. Under certain conditions, both of these behaviors can be observed in some color variants of the fluorescent Sindbis virions, as described in a later section. As a general check for the contributions to the variance from any sort of time-dependent photophysical behavior, images of virions were collected using exposure times ranging from 5 msec to 5 sec, and successive images of the same field of view were obtained with very short or very long exposure times combined with very short or very long (50 msec, 1 minute) intervals between the exposures. Some of these protocols did have small effects on the *average* brightness (see below), but none had any significant effect on the *variance* of measured brightness among fluorescent virus particles.

A clue to the origin of the unexpectedly large variance came from empirical observation of increased variance in images from slides that for some reason had noticeably higher background fluorescence. Normally the background fluorescence in images of the fluorescent Sindbis is very low, of the order of 1% of the fluorescence of a virus particle. Thus I had assumed that the very low background could not contribute significantly to the observed large variation in virus particle brightness. Nevertheless, model calculations were done to evaluate this assumption, as well as to investigate the effect of various forms of intrinsic virus heterogeneity on the brightness measurements. As shown in Figure 7, the background fluorescence, more specifically, the variance of the background fluorescence, does in fact have a profound effect on the width of the histogram of net brightness measured in images of a population of randomly located virus particles, even though the average background is very dim relative to the virions. The *average* magnitude of the background fluorescence relative to the virus particle has only a small influence on the width of the distribution if the background is relatively smooth. However, spatial *variability* in the background has a dramatic effect, even though the full range of background fluorescence is a fraction of one percent of the peak fluorescence from the virions (Figure 7).

**Figure 7.**
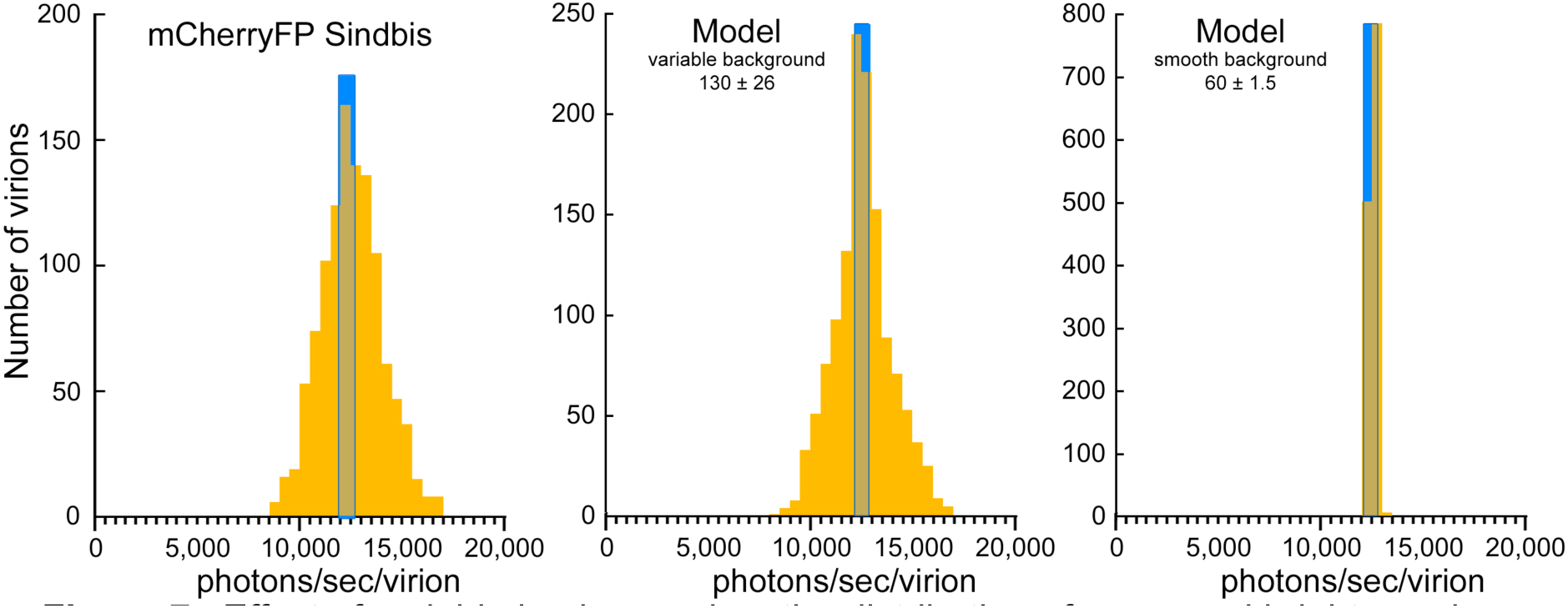
Effect of variable background on the distribution of measured brightness in images of a population of identical fluorescent virus particles. *Left*, experimental data; *Middle*, simulation with average background set to 1% and standard deviation of 0.2%, relative to peak virus brightness; *Right*, simulation with average background 0.5%, standard deviation 0.01%. In all three histograms the blue bar corresponds to the expected full width of the distribution based solely on Poisson noise in the fluorescence signal from a virus particle.

Experience gained with quantitative analysis of virus brightness led to the realization that measurements of virus brightness using the same sample and same microscope varied over time. Likewise, measurements of the same sample on the same day but using two different microscopes usually gave quite different results. As one would expect, the primary cause of these differences is the variation over time or between microscopes in the intensity of illumination used to excite fluorescence. A simple way to avoid this problem is provided by the nearly constant ratio of quantum yield for fluorescence relative to quantum yield(s) for “photobleaching” for each FP. As long as that ratio is constant, then the average number of photons of fluorescence emitted by each FP molecule before it is destroyed by photobleaching, or driven into a long-lived dark state, will also be constant. (In what follows, “photobleaching” with quotation marks will be used as a catch-all term for both the irreversible light-induced destruction of the fluorophore as well as light-induced driving of the fluorophore into any dark state from which recovery is slow compared to the duration of the time-lapse image series used here, which is of the order of 30 sec.) Of course, for any one individual FP molecule, the actual number of photons emitted before “photobleaching” is unpredictable and highly variable, but for an ensemble of 240 molecules, the total number of photons emitted before, say, half of them become dark has a narrow distribution. The mean value of that distribution would be expected to be independent of illumination intensity. Higher irradiance bleaches or drives molecules into a dark state faster, but in exact proportion it also generates fluorescent photons at a higher rate. Decreasing irradiance likewise decreases “photobleaching” rate and photon emission rate by exactly the same factor. This linkage suggests a more reproducible assay for FP brightness: total photons detected during a series of exposures that results cumulatively in a decrease of brightness to 50% of its initial value, a quantity that will be denoted “Half-Yield” (***HY***). Figure 8 gives an example of the insensitivity of ***HY*** to changes in illumination intensity. As will be shown below, the use of ***HY*** as a measure of brightness does not completely evade all of the potential inaccuracies resulting from the complex photophysics/photochemistry of FP, but for counting molecules it seems the best measure available. A similar measure has been proposed previously as a means of finessing problems due to concentration quenching (“inner filter” effects) in quantitative fluorescence measurements (Hirschfeld, 1976).

**Figure 8.**
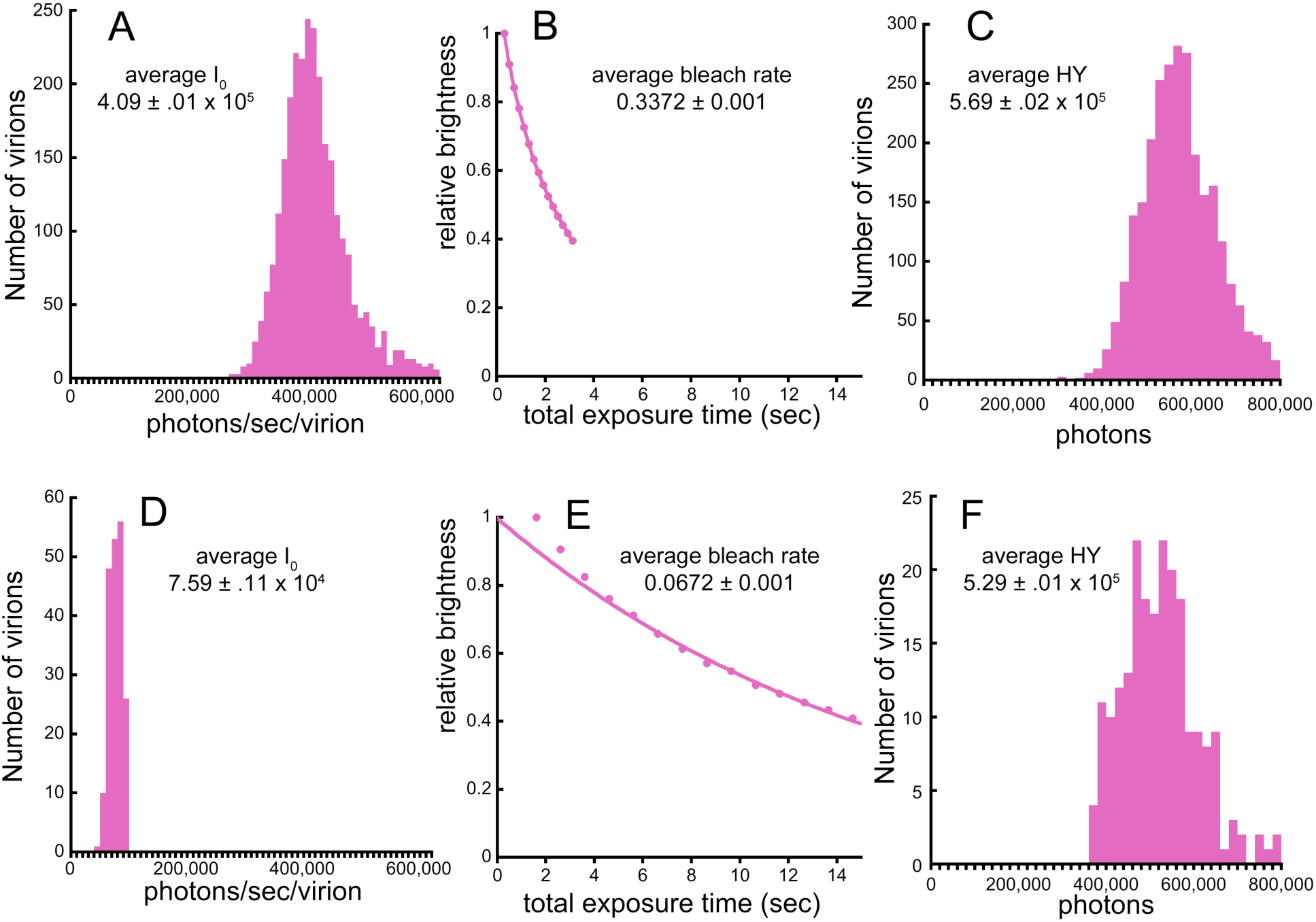
Effect of excitation irradiance on the initial brightness, “photobleaching” rate, and ***HY***. (*A-C*) Illumination ~150 watt/cm^2^; (*D-F*) Illumination ~25 watt/cm^2^. (*A*), (*D*), (*C*), and (*F*) show histograms of measurements on single virus particles. The data points in (*B*) and (*E*) show the average brightness of all particles. The smooth lines in (*B*) and (*E*) are least squares fits to the decay curve between 85% and 50% of initial brightness. Initial brightness (*A vs. D*) and bleach rate (*B vs. E*) are increased by ~5-fold at the higher irradiance, but the total number of photons collected (***HY***) changes by < 10% (C vs. F).

***HY*** can be measured directly by summing the net acquired photons in sequential images of a virus particle or organelle until its brightness falls to half of its initial value. A less noisy estimate of ***HY*** can also be obtained using the ratio of two derived parameters, the initial brightness and the rate constant for “photobleaching”. A formula for the expected value of ***HY*** is obtained by integrating the exponential decay equation, where ***I_0_*** is the initial brightness and ***k*** is the exponential decay constant:

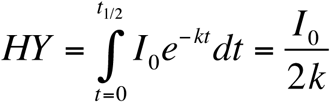

The standard deviation of this estimated sum is related to the standard deviations of ***I_0_*** and ***k*** by:

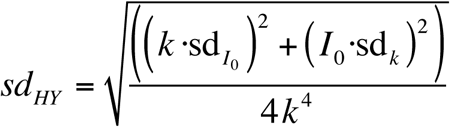

The parameters ***I_0_*** and ***k*** are obtained from photobleaching data. Logarithmic transformation of the equation describing exponential decay of net brightness (***I***) due to photobleaching gives:

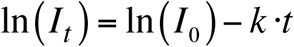

Substituting **y** for ln(***I***), and **x** for **t**, the equation above is put in the standard linear form **y = ax + b.** Estimates of **a** and **b** (i.e., ***k*** and ***I_0_***) are obtained from weighted least squares fitting of a straight line to the transformed equation describing decay. Slightly more accurate estimates are obtained if account is taken of the intrinsic uncertainty in the measured quantities. Due to Poisson noise, the experimentally measured values are known with unequal preciseness, and should thus be weighted differently when used to estimate the parameters ***k*** and ***I_0_***. If the measured quantities are expressed in photons or photons/sec, and assuming Poisson statistics, the variance of ln(***I***) is 1/***I***, so the appropriate weighting factor (***w***) is simply ***I***. With those substitutions and weights, the weighted least squares estimates of the parameters ***k*** and ***I_0_*** are given by (Taylor, 1982; Steel et al., 1997):

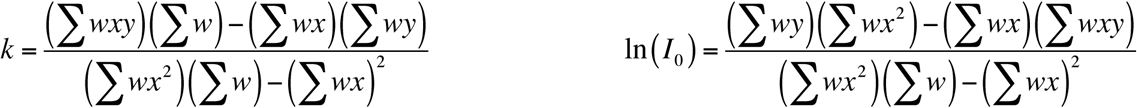

with standard deviations:

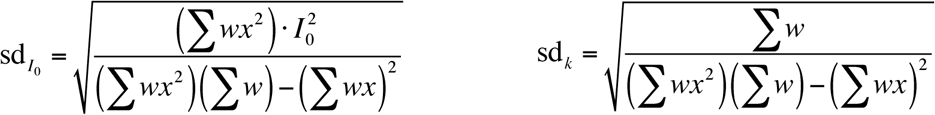

One slight complication is that “photobleaching” curves for many of the fluorescent proteins are not characterized completely by a single exponential over the entire decay. A typical pattern observed in wide-field microscope measurements is a small initial rapid drop in brightness of ~10%, followed by single exponential decay to < 50% of the initial brightness, and then by decay that is slower than predicted by the middle portion of the curve. The first two phases, initial rapid drop followed by single exponential decay, are clearly revealed in Figure 8E. This behavior can be accurately simulated by a 4-state model comprising the normal ground state G0, the excited singlet state S1, and two non-fluorescent dark states, D1 (singlet; accessible from S1 via a first-order transition) and D0 (ground state; accessible from D1 via a first-order transition) (Dean et al., 2011). Both S1 and D1 are subject to “photobleaching”. The rate of recovery from D0 to G0 is slower than the excitation rate of G0 to S1 achieved in a wide-field microscope. The initial rapid drop in brightness corresponds to the decrease of G0 and pile-up of a substantial fraction of the molecules in D0, after which the steady-state rate of “photobleaching” is maintained. For the purpose of counting molecules, the practical consequence of these complications in photophysical behavior is that estimation of the “photobleaching” rate constants needs to be done using only the single-exponential phase of the decay data.

Assayed in this way, by combining the least squares estimates of initial brightness and “photobleaching” rate, measurements of virus brightness in the form of ***HY*** become more reproducible. That reproducibility enabled a systematic characterization of the long term stability of the Sindbis viruses, as well as the brightness under diverse conditions relevant to their use as calibration standards for counting molecules in cells. Table I gives a summary of measurements made over an extended period and in radically different conditions. Note that all images for quantitative analysis were acquired using wide-field fluorescence microscopes. The second of the two measurements for Cerulean3FP Sindbis was made on the same preparation of virus after it had been stored in buffer at 4°C for four months, showing that the virus particles are quite stable. Measurements on mCherry Sindbis show that the virions lose brightness but retain most of their cumulative fluorescence after formaldehyde fixation or embedding in a transparent epoxy resin (Epotek^®^ optical cement). Mounting in a commercial antifade reduces rather than improves cumulative photon yield.

**Table I.**
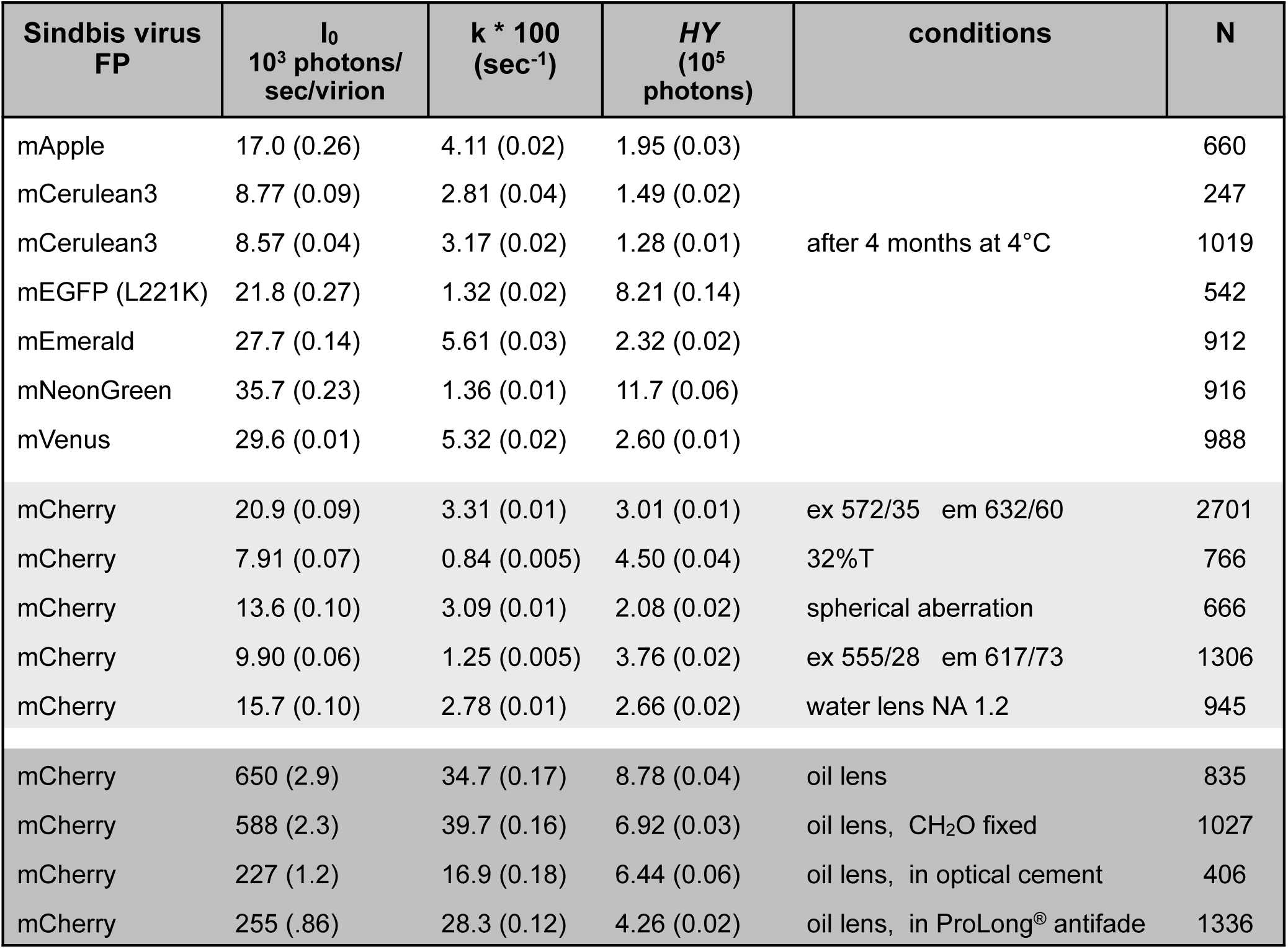
Initial brightness, “photobleaching” rate, and Half-Yield for fluorescent Sindbis virions under various conditions. The values are listed as mean (s.e.m). The data were acquired on 4 different wide-field fluorescence microscopes over a period of 3 years. The five rows of values for mCherryFP Sindbis in the light grey shaded area were all acquired on the same microscope from a single slide on the same day. A silicone immersion lens 60X NA1.3 with correction collar set to minimize spherical aberration was used except as indicated. For one of these mCherryFP Sindbis datasets, the correction collar was deliberately misadjusted to show the effect of spherical aberration, and for another, a water immersion lens 60X NA1.2 was used. The four rows of values for mCherryFP Sindbis in the dark grey shaded area were also acquired together from a single slide on the same day on the same microscope, using an oil immersion lens 60X NA 1.42

Ideally ***HY*** would be completely independent of illumination intensity, and in practice this is often nearly true (Figure 8), but large decreases in intensity do have a measurable effect. An example is given for mCherry Sindbis virions in Table I, where reducing the rate of excitation, either by introducing a neutral density filter with 32% transmittance or by using an excitation bandpass filter with wavelength slightly shifted from the optimum for mCherryFP, increases ***HY*** by 20-50%. The magnitude of this effect is variable with different buffer compositions in the mounting medium. The effect can be explained and quantitatively modeled as a manifestation of a more bleach-resistant singlet state similar to the long-lived dark state previously described for several fluorescent proteins (Sinnecker et al., 2005; Day and Davidson, 2014). Note that this behavior is not predicted by the simple four-state model (Dean et al., 2011) mentioned above. To simulate the effect of illumination intensity, the model must include a transition to a more bleach-resistant fluorescent singlet state with a rate that is *independent* of illumination intensity (i.e., *not* proportional to S1). Fortunately, as shown below, the ***HY*** for FP in Sindbis virions and in cellular organelles seem to be equally prone to these complications in photophysical behavior, so the virions remain useful calibration standards as long as the measurements on virus and cellular organelle are done under similar conditions. Two examples of counting molecules by this method within organelles of the human parasite *Toxoplasma gondii* are described in detail below.

Any parameter affecting the efficiency with which fluorescence emission is detected has an easily measurable effect on ***HY***. For instance, spherical aberration, which smears the image of the fluorescent object along the optical axis, or use of a lower NA objective, which collects a smaller fraction of the emitted fluorescence, both lead to a decrease in the measured ***HY*** (Table I). However, as with illumination intensity, lowered efficiency would not hamper counting the number of FP molecules in an unknown sample if both virions and cellular sample were imaged under the same conditions.

The sensitivity of ***HY*** to detection efficiency means that measurements of the value of ***HY*** for a particular fluorescent Sindbis virus provide a way of quickly comparing the performance of different fluorescent microscopes, similar to but more convenient than the measure reported earlier (Murray et al., 2007).

### Counting molecules of a tubulin-binding protein in a tubulin-based macromolecular assembly

The human pathogen *Toxoplasma gondii* is an obligate intracellular parasite and, like the other members of the Phylum Apicomplexa, contains several specialized organelles that are used for invading host cells. One of these organelles is a cone-shaped assembly, the “conoid”, which is built from 14 spirally arranged fibers that are non-tubular polymers of tubulin, a ubiquitous cytoskeletal protein (Figure 9). The tubulin used to make the conoid fibers also makes canonical microtubules (i.e., hollow tubes) elsewhere in the same cell. The special arrangement of tubulin in the conoid fibers is dictated by non-tubulin components. TgDCX is one such component (Hu et al., 2006; Nagayasu and Murray, 2007). Loss of TgDCX radically disrupts the structure of the conoid and severely impairs invasion of host cells by the parasite (Nagayasu et al., 2016). The non-tubular polymeric form of tubulin in the conoid of the parasite is not found in the cells of their host, suggesting that TgDCX may be an attractive target for new parasite-specific chemotherapeutic agents. Thus in addition to its intrinsic interest as a building block of a stunning piece of cellular architecture, TgDCX and its role in conoid assembly are worthy of study from a clinical standpoint. It would be particularly useful to know the stoichiometry of TgDCX relative to tubulin, so that one could begin to formulate concrete models for how the conoid is constructed.

**Figure 9.**
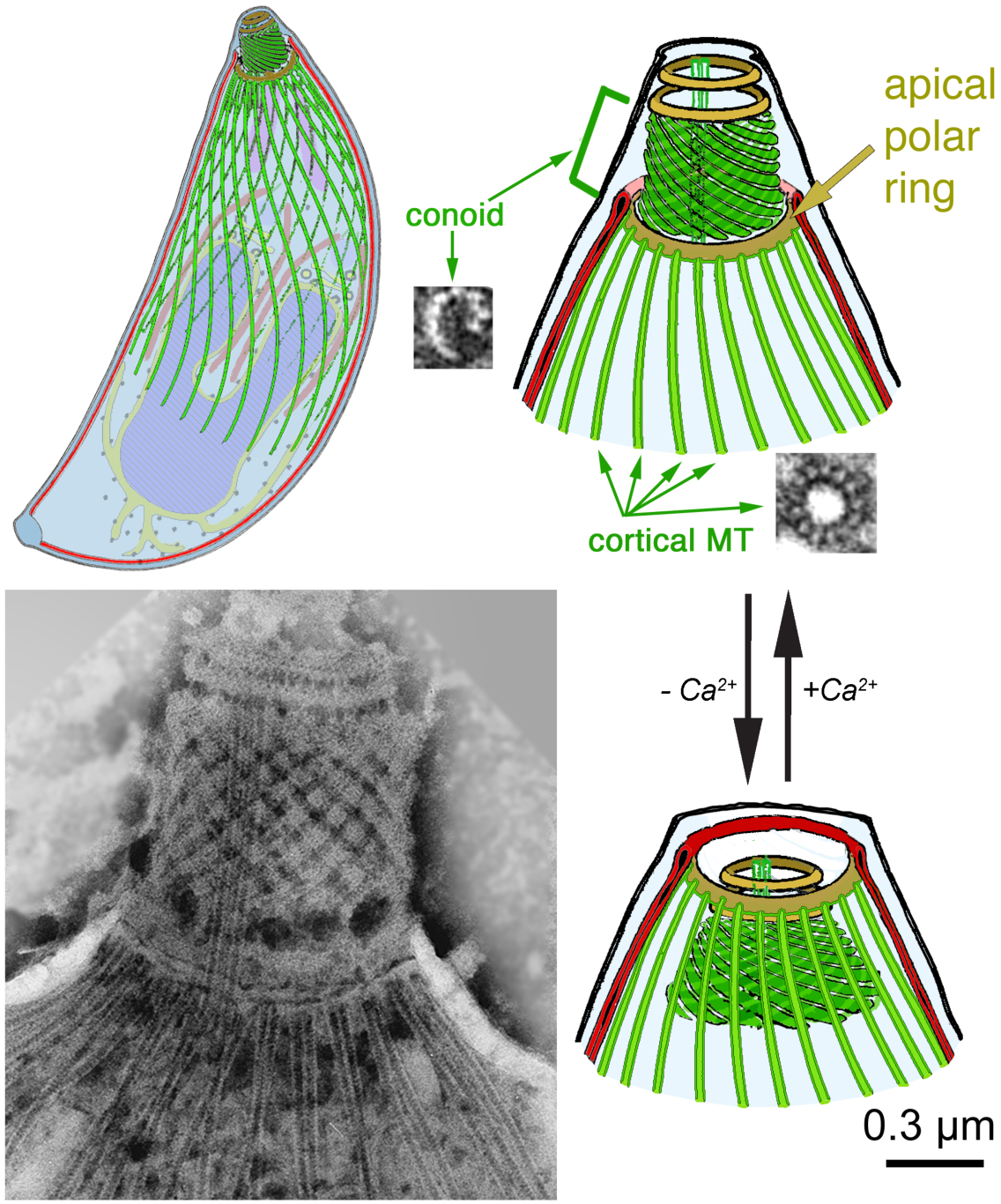
Drawings and EM images of *Toxoplasma gondii* showing various features mentioned in the text. Drawings modified from (Liu et al., 2016). Lower left EM image by Dr. Naomi Morrissette (Morrissette et al., 1997). The small inset EM images show two of the tubulin-containing polymers in *T. gondii*, including the inverted-“J”-shaped conoid fibers that contain TgDCX. The conoid extends and retracts through the apical polar ring as shown, in response to changing [Ca^2+^]. The apical polar ring contains the protein TgAPR1.

Three different Toxoplasma lines were constructed, utilizing mCherryFP-TgDCX (Nagayasu et al., 2016), or TgDCX-mCherryFP, or TgDCX-mNeonGreenFP to replace the endogenous TgDCX by homologous recombination. Figure 10 shows wide-field and structured illumination microscopy (SIM) images of the mCherryFP-TgDCX homologous recombinant parasites. These parasites are all clonal (i.e., all the parasite cells are identical). Their genome contains the FP-tagged copy of TgDCX only (i.e., every molecule of TgDCX in every cell is fused to one mCherryFP molecule). The fluorescence is restricted to the conoid region of the parasite. The results of measurements (s.e.m.) of fluorescence “photobleaching” on cells of these three lines and on the corresponding fluorescent Sindbis virions are listed in Table II. The derived estimates for the number of TgDCX molecules in the conoid of the mCherryFP-TgDCX and TgDCX-mNeonGreenFP lines are identical (2574 ± 20 and 2551 ± 73), but the number derived from the TgDCX-mCherryFP line is significantly different (1740 ± 59).

**Figure 10.**
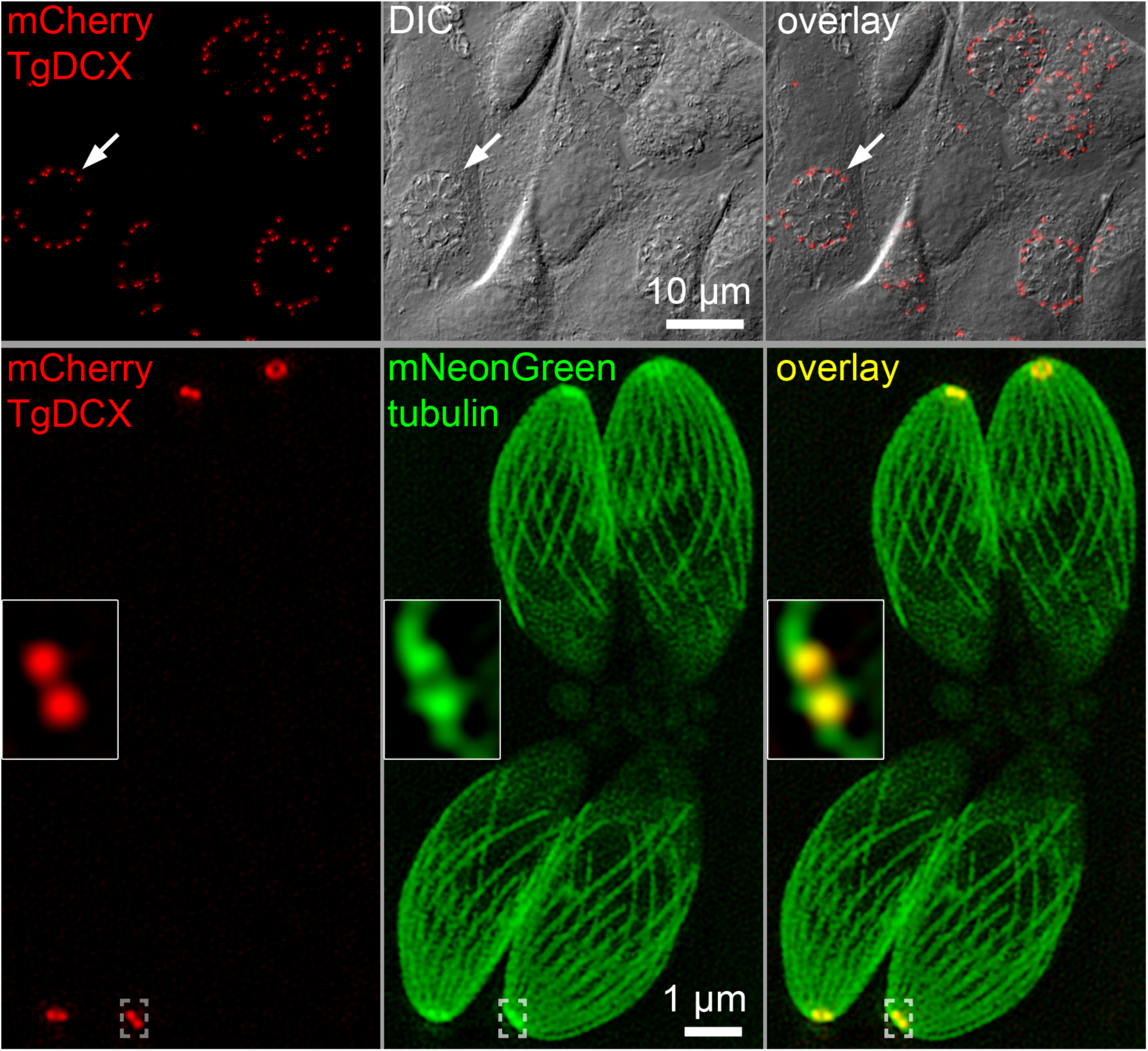
Wide-field epifluorescence, DIC, and SIM images of *T. gondii*. (*upper row*) A monolayer culture of fibroblasts infected with recombinant parasites in which the TgDCX gene has been replaced with DNA coding for an mCherryFP-TgDCX fusion protein. *T. gondii* is an obligate intracellular parasite and lives within a vacuole in the host cell. The white arrow points to a vacuole containing 16 parasites, each of which has one spot of mCherryFP fluorescence at its apical end. The field of view includes four 16-parasite vacuoles plus a few other smaller ones. (*lower row*) SIM images of four mCherryFP-TgDCX knock-in parasites that are also expressing mNeonGreenFP-β1 -tubulin. mNeonGreenFP fluorescence is seen in both the cortical microtubules and conoid, which is retracted as usual in these intracellular parasites (cf Figure 9). mCherryFP fluorescence is restricted to the conoid. The insets show a 4X enlargement of the conoid region of one parasite, as marked by the dashed brackets. In these images, the contrast of the red and green channels has been adjusted independently so that both will be visible in the overlays.

**Table II.** Data for counting TgDCX molecules in *T. gondii*. Values are given as mean (s.e.m). The last column gives the number of FP molecules per conoid or per virus particle.

| | $I_0$<br>(photons/<br>sec) | bleach rate<br>(sec<sup>-1</sup>) | $HY$<br>(photons) | N | number of<br>FP<br>molecules |
| --- | --- | --- | --- | --- | --- |
| mCherryFP-TgDCX | 308,000 (3100) | 0.028 (0.001) | $4.87 (0.04) \times 10^6$ | 361 | 2574 (20) |
| TgDCX-mCherryFP | 133,000 (3400) | 0.019 (0.001) | $3.23 (0.11) \times 10^6$ | 92 | 1740 (59) |
| TgDCX-mNeonGreenFP | 232,000 (6800) | 0.011 (0.003) | $1.24 (0.04) \times 10^7$ | 140 | 2551 (73) |
| mCherryFP Sindbis | 27,600 (60) | 0.027 (0.001) | $4.55 (0.01) \times 10^5$ | 1999 | 240 |
| mNeonGreenFP Sindbis | 35,700 (230) | 0.014 (0.001) | $1.17 (0.06) \times 10^6$ | 916 | 240 |

One possibility is that the discrepancy reflects a real difference in the number of molecules of TgDCX in the conoids of lines expressing mCherryFP fused to the N-terminus versus the C-terminus. However, from a biological standpoint, a difference among the three lines in their conoid content of TgDCX seems unlikely. In the absence of TgDCX the structure of the conoid is severely disrupted and the rate of growth of the parasite is greatly impaired (Nagayasu et al., 2016). In the three knock-in lines, both the structure of the conoid and the rate of growth is normal (*data not shown*), suggesting that the number of molecules of FP-tagged TgDCX in the conoids of all three knock-in lines is the same as TgDCX in wild-type.

Alternatively, the discrepancy in estimated number of TgDCX molecules per conoid might arise from some difference in the photophysical properties of mCherryFP in the conoid when fused to the N-terminus of TgDCX versus the C-terminus of TgDCX — different extinction coefficient (ε), quantum yield for fluorescence (**Φ_*F*_**), or quantum yield for “photobleaching” (**Φ_*B*_**). It would be difficult to measure directly any of those fundamental photophysical parameters individually for mCherryFP in the conoid, but the parameters ***HY***, ***I_0_***, and ***k***, which I have measured, are linearly related to combinations of ε, **Φ_*F*_**, and **Φ_*B*_**. In each case, the proportionality constant includes the excitation photon flux.

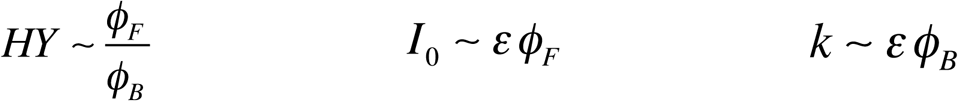

***HY*** should be unaffected by changes in the extinction coefficient. Therefore a difference in ***HY*** between mCherryFP fused to the N-terminus versus the C-terminus of TgDCX means that the ratio of the quantum yields for fluorescence and “photobleaching” must be different for those two fusion proteins. Because both ***I_0_***, and ***k*** are also different for the two (Table II), it seems probable that **Φ_*F*_** and **Φ_*B*_** must both change when mCherryFP is switched from N- to C-terminus. (The measurements for the two lines were made under identical illumination conditions.) Other observations (see below) also suggest that **Φ_*F*_** and **Φ_*B*_** are sensitive to their local environment. With this sensitivity in mind, it seems prudent to ensure that the measured “photobleaching” rates are the same for the virion and the cellular organelle before using the fluorescent virus, or any other form of FP, as a calibration.

### Counting molecules of a protein component of a putative microtubule organizing center

The apical polar ring, from which the 22 cortical microtubules (MTs) of *T. gondii* originate, is a distinguishing feature of all apicomplexans (Morrissette and Sibley, 2002) and is hypothesized to be the primary MT organizing center for the *Toxoplasma* cytoskeleton (Russell and Burns, 1984). TgAPR1, a 460 aa protein, has been identified as a component of the apical polar ring (Hu et al., 2006; Zhang and Murray, 2006). A line of knock-in parasites was constructed in which the single genomic copy of TgAPR1 was replaced with TgAPR1-mCherryFP (Figure 11) by homologous recombination (Wu et al., 2016).

**Figure 11.**
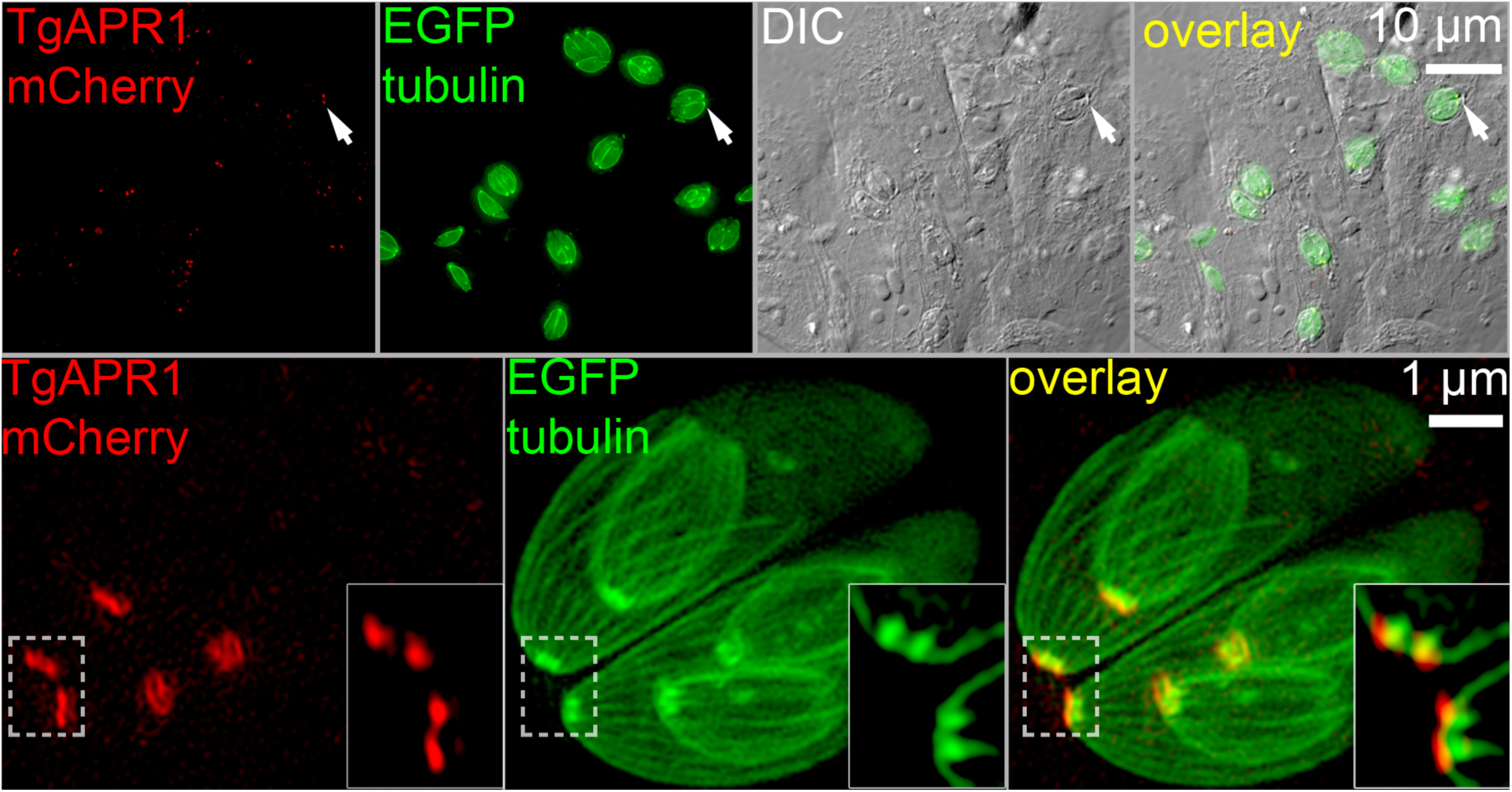
Wide-field epifluorescence, DIC, and SIM images of *T. gondii*. (*upper row*) A monolayer culture of fibroblasts infected with recombinant parasites in which the TgAPR1 gene has been replaced with DNA coding for a TgAPR1-mCherryFP fusion protein. The parasites also contain an extra copy of the gene for β1-tubulin, fused to EGFP. The white arrows point to one of the many vacuoles containing 2 parasites. (*lower row*) SIM images of TgAPR1-mCherryFP knock-in parasites which are also expressing EGFP-β-tubulin. EGFP fluorescence is seen in both the cortical microtubules and the conoid, which is retracted in these parasites. mCherryFP fluorescence is restricted to the apical polar ring, which surrounds the bright spot of EGFP tubulin in the conoid (cf Figures 9 and 10). The insets show a 4X enlargement of the apical polar ring regions of two parasites, as marked by the dashed white border. In these images, the contrast of the red and green channels has been adjusted independently so that both are visible in the overlays.

“Photobleaching” of the fluorescent ring structure in the knock-in parasites was measured along with mCherryFP-Sindbis virus under the same conditions. From the ***HY*** of the ring structure and the virus (2.37 ± 0.04 × 10^6^ and 4.55 ± 0.01 × 10^5^ photons respectively), the number of TgAPR1 molecules in the ring was calculated to be 1251 ± 21. The bleach rate for mCherryFP in the ring was the same as for the mCherryFP-Sindbis, 0.027 sec^−1^.

An interesting phenomenon was observed with this parasite line when “photobleaching” measurements were carried out on the parasites outside their host cell, suspended in buffer under a coverslip. In this situation, two distinct “photobleaching” behaviors were observed. In some of the extracellular parasites, the TgAPR1-mCherryFP fluorescence decayed much faster, ~10-fold faster, than it did in parasites located within the host cell. Intermingled among this fast-bleaching population were other parasites whose TgAPR1-mCherryFP fluorescence decayed at about the same rate as all intracellular parasites, which was the same as the decay rate of the mCherryFP-Sindbis virus. Close inspection of DIC images of the extracellular parasites revealed that the slow-bleaching population were lysed, as judged by the presence of Brownian motion of small particles in the cytoplasm. The fluorescence of the fast-fading population recovered substantially over a period of 10−20 minutes. Recovery of the slow-fading population was much less complete. In contrast, if the parasites were in an open dish, covered with a thin layer of buffer but not covered by a coverslip, the fast-bleaching population was not observed. Extracellular parasites remain alive and respiring for hours, so oxygen is likely to be soon depleted when they are mounted in a tiny volume between coverslip and slide.

Presumably the difference between the lysed and intact extracellular parasites is related to some metabolism-dependent change in the parasites when they are outside the host and metabolically stressed. A few experiments were undertaken to confirm that the external chemical environment could significantly affect the rate of loss of mCherryFP fluorescence. Methylene blue is a phenothiazine dye that generates reactive oxygen species when illuminated (Tardivo et al., 2005). Adding 20 μM methylene blue to the culture medium in which the extracellular parasites were suspended eliminated the difference between the two populations. In the presence of methylene blue, TgAPR1-mCherryFP fluorescence in the ring structure decayed at a fast rate in all parasites, lysed or intact, and showed significant recovery after 10-20 minutes. mCherryFP in the Sindbis virus was affected similarly, “bleaching” fast, but with significant recovery after 10-20 min. On the other hand, scavengers of oxygen (glucose plus glucose oxidase plus catalase) or superoxide (ascorbic acid plus superoxide dismutase) had little effect on the rate of fading of either the fast or slow population.

## Methods

### Plasmid construction

#### TE12-mCherryFP

A plasmid (TE12) encoding the complete genome of recombinant Sindbis virus (Lustig et al., 1988) was kindly provided by Dr. Suchetana Mukhopadhyay (Indiana University, Bloomington). From TE12 a 3462 bp region coding for proteins C, E3, E2, 6K, and part of E1 was removed with BsiWI and HpaI. The gap was filled with a 4206 bp piece containing the same coding sequences plus mCherryFP fused in-frame between E3 and E2. Overlap PCR from three fragments was used to produce the 4206 bp piece. The templates used for PCR, the primers, and the lengths of the three fragments are: Fragment #1 (1758 bp) amplified from the TE12 plasmid, position 6910-8630; primers S1 and AS2 (Table III). Fragment #2 (785 bp), from a mCherryFP coding sequence (synthesized with silent mutagenesis to remove multiple restriction sites); primers S2 and AS5. Fragment #3 (1804 bp), from TE12 position 8629-10396; primers S5 and AS7. The three purified fragments were mixed and ligated by PCR with primers S1 and AS7. The 4206 bp product, cut with BsiWI and HpaI, was ligated into the cut TE12 vector. The furin recognition site at the C-terminus of E3 (SGRSKR) is intact in the expressed protein product. Before cleavage by furin, the N-terminus of mCherryFP is separated from the furin cleavage site by a 5 aa flexible linker GAPGSA. The C-terminus of mCherryFP is coupled to the N-terminus of E2 via the 6 aa linker AAAGSG.

**Table III.**
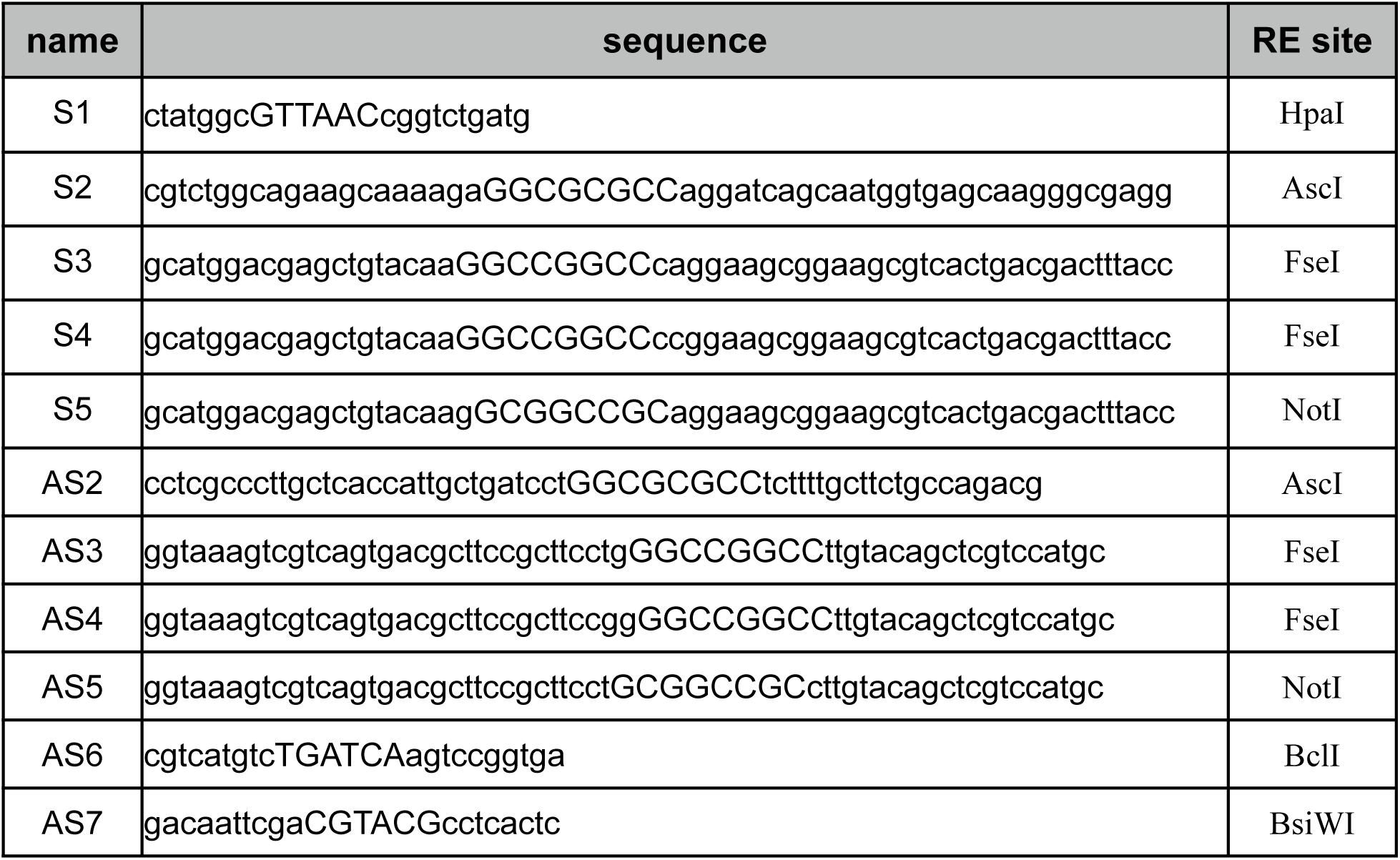
Primers used in this project. Restriction enzyme recognition sequences are capitalized.

#### TE12-mAppleFP, TE12-mEmeraldFP, TE12-mNeonGreenFP, TE12-mVenusFP

These four were prepared in the same way. In each case, fragment #2 for the overlap PCR was obtained by amplifying the corresponding FP coding sequence with primers S2 and AS5. The plasmid template for mNeonGreenFP (Shaner et al., 2013) was kindly provided by Dr. Richard Day (Indiana University School of Medicine, Indianapolis).

#### TE12-EGFP

The region of TE12 coding for proteins C, E3, E2, 6K, and part of E1 (3267 bp) was excised with BsiWI and BspQI and replaced with a 4020 bp piece containing the same coding sequences plus EGFP fused in frame between E3 and E2. The 4020 bp piece was cut from a 4240 bp piece produced by overlap PCR from three fragments. Fragment #1 (1758 bp), from TE12, position 6910#8630; primers S1 and AS2 (Table III). Fragment #2 (794 bp), from an EGFP-containing plasmid; primers S2 and AS4. Fragment #3 (1804 bp), from TE12 position 8629-10396; primers S4 and AS7. These three fragments were purified, mixed, and used as template for PCR with primers S1 and AS7.

#### TE12-mCerulean3FP

A 2244 bp region of plasmid TE12 coding for proteins C, E3, and part of E2 was removed by cutting with BclI and BspQI, and replaced with a 2997 bp piece containing the same coding sequences plus mCerulean3FP fused in frame between E3 and E2. The 2997 bp piece was cut from a 3216 bp piece produced by overlap PCR from three fragments. Fragment #1 (1758 bp) was amplified from TE12 position 6910-8630 with primers S1 and AS2 (Table III). Fragment #2 (794 bp) was amplified from plasmid pmCerulean3FP-N1 ((Markwardt et al., 2011) kindly provided by Dr. Richard Day) with primers S2 and AS3. Fragment #3 (780 bp) was amplified from TE12 position 8629-9372 with primers S3 and AS6. These three fragments were purified, mixed, and used as template for PCR with primers S1 and AS6.

#### TE12-tdTomatoFP

The EGFP CDS of TE12-EGFP was removed with enzymes AscI and FseI, and replaced with a sequence coding for tdTomatoFP, prepared by PCR amplification with primers S2 and AS4. The linker between the C-terminus of E3 and N-terminus of the FP, which includes an AscI site, codes for the amino acid sequence GAPGSA. Between the C-terminus of the FP and N-terminus of E2, the linker codes for the amino acid sequence AGPGSG and includes an FseI site.

#### TE12-mEGFP(L221K)

A 4215 bp region of plasmid TE12-mVenusFP was removed by cutting with BsiWI and HpaI, and replaced with a 4020 bp piece containing the same coding sequences with mEGFP(L221K) instead of mVenusFP fused in frame between E3 and E2. The 4020 bp piece was produced by overlap PCR from three fragments. Fragment #1 (1758 bp) was amplified from TE12-mCherryFP position 6910−8667 with primers S1 and AS2 (Table III). Fragment #2 (794 bp) was amplified from an mEGFP(L221K)-containing plasmid (Addgene #21042) with primers S2 and AS5. Fragment #3 (1804 bp) was amplified from TE12-mCherryFP position 9337-11140 with primers S5 and AS7. These three fragments were purified, mixed, and used as template for PCR with primers S1 and AS7. In TE12-mEGFP(L221K) the E3-FP linker is GAPGSA and the FP-E2 linker codes for AAAGSG and includes a NotI site.

#### In vitro transcription

Sindbis viral RNA (+)-strand was transcribed *in vitro* from the TE12-FP plasmids with SP6 RNA polymerase (Rice et al., 1987). PvuI-linearized plasmid DNA (100-200 ng) was precipitated with ethanol, re-dissolved in 20 μL of 40 mM TrisCl (pH 7.9 at room temperature), 10 mM MgCl2, 2 mM spermidine, 10 mM DTT, 2 mM each of rATP, rCTP, rGTP, and rUTP, 0.5 mM 3´-O-Me-m^7^G(5’)ppp(5’)G RNA cap structure analog (New England Biolabs #S1411),1X RNAsecure™ reagent (Life Technologies #AM7005), and incubated at 60°C for 10 min. After cooling to room temperature, 20 units of SUPERase• In™ RNase inhibitor and 20 units of SP6 RNA polymerase (New England Biolabs #M0207) were added and the mixture was incubated at 39°C for 2 hours. The entire reaction mix was immediately used for transfection.

### RNA transfection and virus production

Each *in vitro* transcribed RNA was transfected into Vero or BHK-21 cells in a 24-well plate using Effectene (Qiagen #301425) following the manufacturer’s protocol, with a total volume of 180 μL of serum free media per well. After overnight incubation at 37°C in a CO_2_ incubator, 0.8 mL of culture medium was added and incubation continued for an additional 24 hours. By 6-8 hours after transfection, plaques of virus infected cells were easily visible by fluorescence microscopy using a low power (4x or 10x) objective. 24-48 hours after transfection, the culture supernatant was removed, centrifuged at 16,000 x g for 5 min to remove cell debris, then stored in 0.1 mL aliquots frozen at -80°C. The concentration of infective virus particles was determined by plaque assay (Hernandez et al., 2010). As is typical of even wild-type Sindbis (Knight et al., 2009), the number of infective particles measured by plaque assay is much smaller, 1% or less, than the number of virus particles determined by physical or optical measurements. With newly prepared virus stocks, 100% of the plaques are fluorescent, but after 5−6 serial passages, dark mutants appear and seem to have a growth advantage, as the fraction of plaques that are fluorescent declines steadily with each additional passage.

### Virus production and purification

T150-flasks of Vero or BHK-21 cells were incubated with Sindbis-FP virus, ~0.2 pfu/cell, diluted from frozen stock into a volume of culture medium (DMEM, 2 mM glutamine + 12% (Vero) or 2% (BHK) calf serum (HyClone™ Cosmic Calf™ Serum, Thermo Scientific #SH30087) just sufficient to cover the cells and rocked at room temperature for 1 hour. 20 mL culture medium was added and the flasks were incubated at 37°C for 24−48 hr. The culture supernatant was removed, centrifuged at 5000 x g for 20 min, and passed through a 0.22 μm filter. In most cases, this clarified culture supernatant can be used directly as the source of fluorescent virus particles for molecular counting. When further purification was needed (e.g., in attempts to reduce background fluorescence), solid NaCl and PEG-8000 (Sigma-Aldrich #P5413) were added to give final concentrations of 1.0 M and 10% (wt/ vol), and the mixture was stirred gently at room temperature until the solids were dissolved, then at 4°C overnight. Precipitated virus was collected by centrifugation at 20,000 x g for 30 min. The pellet was gently re-dissolved in 1.2 mL buffer H (50 mM Na-HEPES pH 7.0, 0.1 M NaCl, 0.5 mM Na2EDTA) and centrifuged at 16,000 x g for 10 min. The supernatant was removed and saved, and the pellet was extracted with 0.5 mL buffer H and centrifuged at 16,000 x g for 10 min. The combined supernatants were layered over a discontinuous sucrose gradient (1 mL 65% sucrose in buffer H, 2 mL 20% sucrose in buffer H) and centrifuged at 240,000 x g in a Beckman SW55Ti rotor for 45 min. The strongly colored band of purified virus at the 20%−65% sucrose interface was removed with a syringe and 26G needle, diluted 5 fold with buffer H, aliquoted and stored frozen at -80°C. The level of fluorescent background can be reduced by this purification, but in my hands it invariably led to an increase in the number of dimers and larger aggregates.

### Biological properties and handling of the Sindbis virus standards

Sindbis virus is a member of the alphavirus family, which includes some important human pathogens (Lloyd, 2009). In the wild, Sindbis itself is an arthropod borne vector spread by mosquitos with a host range spanning birds and mammals, including primates. In humans, particularly in Europe, Africa, the Middle East, and Russia, Sindbis infections are fairly common. Most infections are probably asymptomatic, with the remainder typically involving a self-limited bout of mild fever and arthralgia or skin rash. Human to human transmission does not occur (Schmaljohn and McClain, 1996). Sindbis is categorized as a Risk Group 2 agent and subject to BSL-2 precautions for laboratory work. It is rapidly inactivated by simple disinfectants, including 70% ethanol and the detergents normally used for hand-washing. From the experience of many years of research in many laboratories, the common laboratory strains of Sindbis virus have not proven hazardous to humans. No case of laboratory acquired infection has been reported (PHAC-Ca, 2014).

The quantity of Sindbis virus needed for use as a fluorescent standard is minuscule compared to what is readily generated in the laboratory in a single small culture vessel. Viral suspensions can be stored frozen for years, and new preparations made from frozen stock with minimal effort. For long term use, it is best to make a stock and freeze it, removing small aliquots as needed. Repeated serial passage in culture is not recommended, as dark, faster replicating, mutants appear spontaneously and take over the population.

### Imaging

20 μL of diluted virus culture supernatant or sucrose-gradient purified virus was spread on a cleaned 22x22 mm #1.5 coverslip, allowed to adsorb for a few min, rinsed with buffer, then placed on a 3 μL droplet of mounting medium (Buffer H unless otherwise specified) on a cleaned glass slide. The edges of the coverslip were sealed with VALAP (Vaseline:Lanolin:Paraffin wax 1:1:1). Alternatively, ~500 μL of highly diluted virus suspension was placed in a 35-mm plastic dish with #1.5 glass coverslip bottom (MatTek P35G-1.5–10-C), allowed to absorb for 15-20 min, rinsed with buffer, and covered with 2 mL of buffer. For some measurements, coverslips with adsorbed virus were rinsed, blotted on filter paper, immediately placed on a small droplet of optical cement (Epotek^®^ 305, Epoxy Technology Inc.) on a glass slide and pressed flat, then allowed to harden overnight.

*Toxoplasma gondii* cultures for imaging were grown in a monolayer of human fibroblasts in a MatTek dish. Just before transferring to the microscope, the medium was replaced with CO_2_-independent medium (Gibco 18045–088). A humidified environmental chamber surrounding the microscope stage maintained the sample at 37ºC. For imaging of *T. gondii* outside the host cell, a concentrated suspension of parasites was allowed to adhere to a Cell-Tak^®^ (Corning #354240) coated coverslip or MatTek dish for 20 min in a humid chamber, rinsed with buffer, and mounted as above.

Images for quantitative analysis were acquired on four different wide-field fluorescence microscope systems (three Olympus, one Nikon), using three different types of light sources, ten different filter sets, four different objective lenses, and three different cameras. The various hardware combinations are summarized in Table IV.

**Table IV.**
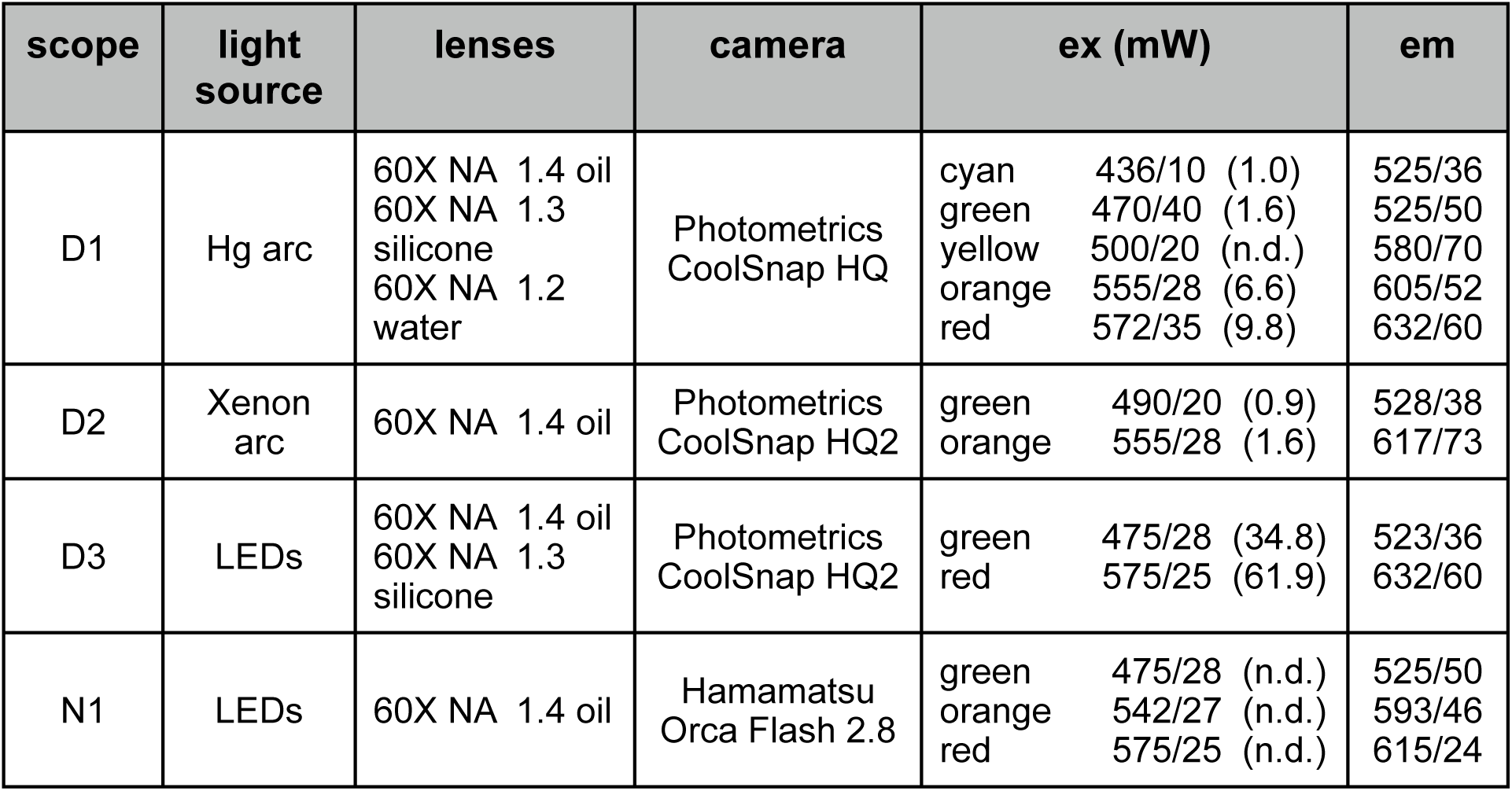
Microscope light sources, lenses, cameras, and filters. D1-3 are different Olympus microscopes associated with different DeltaVision systems. All four microscopes are wide-field. N1 is a Nikon microscope. Excitation power was measured at the objective lens. (n.d.) = not determined.

| scope | light source | lenses | camera | ex (mW) |  |  | em |
| --- | --- | --- | --- | --- | --- | --- | --- |
| D1 | Hg arc | 60X NA 1.4 oil<br>60X NA 1.3<br>silicone<br>60X NA 1.2<br>water | Photometrics<br>CoolSnap HQ | cyan<br>green<br>yellow<br>orange<br>red | 436/10 (1.0)<br>470/40 (1.6)<br>500/20 (n.d.)<br>555/28 (6.6)<br>572/35 (9.8) |  | 525/36<br>525/50<br>580/70<br>605/52<br>632/60 |
| D2 | Xenon arc | 60X NA 1.4 oil | Photometrics<br>CoolSnap HQ2 | green<br>orange | 490/20 (0.9)<br>555/28 (1.6) |  | 528/38<br>617/73 |
| D3 | LEDs | 60X NA 1.4 oil<br>60X NA 1.3<br>silicone | Photometrics<br>CoolSnap HQ2 | green<br>red | 475/28 (34.8)<br>575/25 (61.9) |  | 523/36<br>632/60 |
| N1 | LEDs | 60X NA 1.4 oil | Hamamatsu<br>Orca Flash 2.8 | green<br>orange<br>red | 475/28 (n.d.)<br>542/27 (n.d.)<br>575/25 (n.d.) |  | 525/50<br>593/46<br>615/24 |

### Image analysis

An updated version of the Semper (Saxton et al., 1979) software package (source code generously provided by Dr. Owen Saxton) running under Unix on a Mac Mini or MacBookPro was used for quantitative image analysis. Non-uniform illumination was corrected by “flatfielding” (i.e., dividing each image by the image of a uniformly fluorescent specimen that had been normalized to 1.0 at its peak), and the region for analysis was restricted to the area of the image whose illumination was at least 85% of the maximum. Photosensor measurement of actual illumination intensity during each individual image acquisition, utilizing hardware and software built in to the Applied Precision Deltavision system, was used to globally normalize all images of a dataset to a common standard exposure. Single optical sections at optimal focus were used for analysis of the images, but 3D stacks of 3-5 slices were sometimes used to find the best focal plane. Image brightness in grey-levels was converted to brightness in photons/sec, using an experimentally determined ADU gain calibration (Faruqi et al., 1999; Murray, 2013) together with the known exposure time.

The most troublesome problem to be solved in making quantitative measurements of fluorescence proved to be obtaining an accurate estimate of local background fluorescence. Many different protocols were tried before settling on the following as the most reliable for these particular images. The images considered here are characterized by well-defined small bright spots (virus particles, PSF size) surrounded by large areas of much lower brightness (background). For each image, an under-sampled array of local background values was determined by finding the statistical mode of the population of pixel values in small neighborhoods. The size of the local neighborhoods was typically 0.1% of the area of the full-size image. A full-size background image was then created by bilinear interpolation between the array of local modes.

Images of fluorescent virus normally included >500 virions per field of view. Standard particle analysis algorithms in Semper were efficient at automatic identification of virus particles. The automatic particle recognition algorithm identified patches of brightness satisfying four selection criteria: (1) a contiguous set of pixels with minimum brightness exceeding the local background by a certain threshold, usually taken as >50 times the expected standard deviation of the average background assuming Poisson statistics, and maximum brightness less than four times the predetermined typical peak brightness of a single virus particle; (2) the area of the contiguous patch was required to lie between a minimum (one-half of the PSF area) and maximum (two times the PSF area) size; (3) the patch contained no holes; and (4) the circularity of the patch (defined as area multiplied by 4π and divided by perimeter-squared) exceeded 0.85. The locations of the centers of mass of the identified particles were computed, and particles that had neighbors closer than twice the PSF diameter were excluded from the analysis. For the remaining particles, values of all pixels within an area equal to twice the area of the PSF (calculated for the objective lens and emission wavelength of the FP) centered on the center of mass were summed. The local background value multiplied by the number of pixels within the area of summation was subtracted to give the net brightness for each virus particle.

The raw output from the automated particle recognition programs needs curation before use, as inevitably some objects are picked that satisfy all of the selection criteria but are not single virus particles. To eliminate these aggregates or other spurious objects, filters were applied to the list of candidates output from the particle recognition programs. Fluorescent particles of debris generally behave differently from the virus particles and were efficiently eliminated by filters based on the extent of photobleaching. The exposure times and number of images in a photobleach series were chosen to give ~ 50% bleaching of virus particles. The acceptable window for the filter was 30% < cumulative photobleaching < 70%. The correlation coefficient obtained from fitting a single exponential to the decay curve was also constrained: acceptable corr. coeff. > 0.85. Unresolved aggregates of 2 or more virus particles were eliminated by setting upper limits on the allowed maximum single-pixel brightness within a spot, and the maximum net initial brightness (***I_0_***) of the particle, as calculated from the least squares fit to the decay curve. Appropriate thresholds for discriminating between single virions and unresolved multimers were obvious upon examination of histograms of maximum brightness and ***I_0_*** (e.g., the histograms for mCherryFP-Sindbis and mEmeraldFP-Sindbis in Figure 3).

Images of organelles within *Toxoplasma* contained at most a few dozen organelles per field of view. For this small number, and due to the more variable background, it was simpler to identify organelles manually in displayed images. Subsequent processing, including the estimation of local background, was the same as for the virus particles. The area of summation was set individually for each manually identified organelle by using the cursor to define the major and minor semi-axes of an elliptical ROI. Frequently the images of two organelles belonging to two closely adjacent *Toxoplasma* overlapped, but this situation was readily recognized from the DIC image that was taken along with every fluorescent image, so the measured fluorescence could be reliably annotated as arising from one or two organelles. Using the fluorescence/DIC overlay image it was also straightforward to distinguish organelles belonging to forming daughter parasites from those of the mature adult.

### Modeling of fluorescent virion images and noise analysis

Simulated images of fluorescent virus particles were created as test objects to validate the image analysis programs and to analyze quantitatively the contributions of various sources of noise. Each digital test object was the sum of three components: camera offset plus read-out noise, background fluorescence, and virus particles.

Camera offset was a constant value. Camera read-out noise was modeled as a Gaussian distribution with a mean of zero and a standard deviation of 8 gray levels, which is the experimentally measured value for the camera used for most of the experiments. A 2D array of offset plus read-out noise was created.

Background fluorescence was modeled as the sum of a baseline fluorescence plus a spatially varying but temporally constant component whose amplitude relative to the baseline could be set as a parameter in the program. A 2D array was created, apodized according to the non-uniform illumination function (see below), and Poisson noise corresponding to the mean background fluorescence was added.

Virus particles were modeled as delta functions. The possibility of intrinsic heterogeneity in the virus particles was allowed for in two ways. First, the magnitude of each delta function was chosen by random drawing from a population with Gaussian distributed magnitudes. The standard deviation of the Gaussian distribution was set as a parameter in the program. Second, each virus particle was assigned its own individual photobleaching rate constant, again drawn randomly from a Gaussian distributed population of rate constants with standard deviation as a program parameter.

A series of simulated images modeling successive time-points in a bleach series was created from these three components as follows. First, a fixed number of virus particles (i.e., delta functions, typically 250 to 1000) was distributed randomly over a 2D array. In successive images (i.e., successive time points in the bleach series), the locations of the virus particles were unchanged, but the magnitude of each delta function in the 2D array was decremented exponentially according to its individual photobleaching rate constant. Non-uniform illumination across the field of view was simulated by tapering both the delta function magnitudes and the photobleaching rate constants according to an apodizing function with circular symmetry, with its symmetry axis slightly offset from the center of the 2D array. The apodizing function varied from 1.0 at the center of symmetry to a parameter-defined fraction (typically 0.75) at the periphery of the 2D array. The 2D array of delta functions was then blurred by convolving with a PSF, modeled as a 2D Gaussian with standard deviation of 0.10 µm, appropriate for an imaging system with effective NA of 1.3 and wavelength 632 nm (PSF radius 0.29 µm). The pixel size was set to be 0.11 µm, slightly finer than Nyquist sampling, and corresponding to the actual pixel size in most images. Small random defocus errors, uncorrelated between successive images of the bleach series, were modeled by varying the width of the PSF randomly over a small range, typically by an amount corresponding to a maximum defocus error of 0.15 µm. Finally Poisson noise corresponding to the mean brightness among virus particles was added to the 2D array of blurred delta functions, and the background and camera offset/readout noise arrays were added.

Five to ten simulated bleach series of 15 to 50 time points each were created for each test dataset, comprising 1000 to 10,000 virus particles. Program parameters were held constant across bleach series within a set, but a different spatially modulated background and a different set of random x-y locations of virus particles across the 2D array was generated for each bleach series in the set. The simulated datasets were analyzed with the same suite of image analysis programs as used for the real data.

### Modeling of mCherryFP dark states and photobleaching

The photochemical transitions in mCherryFP under different illumination conditions were modeled by numerical integration of the set of linked differential equations using the COPASI software for biochemical systems simulation (http://copasi.org/). The basic 4-state model, taken from (Dean et al., 2011), included the normal ground state G0, the excited singlet state S1, and two non-fluorescent dark states, D0 and D1. At the rate of excitation achieved in a wide-field microscope, the recovery from D0 to G0 is rate limiting. Both S1 and D1 are subject to irreversible photobleaching. The 5-state model incorporated another fluorescent singlet state that is more stable against photobleaching than S1, accessible from S1 via a pathway with fixed and limited flux. At low rates of excitation, most of S1 enters the more photostable state and is then protected from photobleaching, but as the excitation rate increases, the fixed-rate transition to the photostable state cannot keep up, leading to increased photobleaching of the FP.

### Toxoplasma gondii culture and transformation

*T. gondii* tachyzoites were grown in monolayers of human foreskin fibroblast (HFF) cells (Roos et al., 1994), harvested from culture supernatant by centrifugation at ~1500 x g for 5 min, passed through a 3 μm Nuclepore filter (Whatman 110612), centrifuged, and resuspended in CO2 Independent Medium (Gibco 18045–088). Transfection was carried out as previously described (Hu et al., 2006). For tagging endogenous genes with FPs by homologous replacement, the loxP/Cre recombinase “knock-in” plasmid pTKO-II and protocol (Heaslip et al., 2010) were employed, using a *RH*Δ*ku80*Δ*hx* parasite strain (Fox et al., 2009; Huynh and Carruthers, 2009). Plasmids pTKO4-TgDCX-mNeonGreenFP and pTKO4-TgDCX-mCherryFP, used for homologous replacement of TgDCX by TgDCX-mNeonGreenFP and TgDCX-mCherryFP respectively, were constructed as previously described for plasmid pTKO4-mCherryFP-TgDCX (Nagayasu et al., 2016). The knock-in lines of *T. gondii* expressing TgDCX-mNeonGreenFP and TgDCX-mCherryFP were constructed as previously described (Nagayasu et al., 2016) for the mCherryFP-TgDCX knock-in line.

## Discussion

In this paper I report the construction and characterization of a set of standards for calibrating quantitative measurements of fluorescence arising from cellular macromolecular assemblies tagged with any of seven different fluorescent proteins. These virus-based reagents have a number of advantages over other proposed standards, including ease of preparation, low cost, indefinite lifetime, long-term reproducibility, and straightforward extension to new FPs as they become available. Two applications of the standards to counting molecules in the human parasite *Toxoplasma gondii* were described. As an aid to the effective use of these or other fluorescent standards, I have also highlighted the potential complications and sources of inaccuracy and impreciseness that users are likely to encounter.

### Factors affecting the accuracy and preciseness of counting FP molecules in cells

Some of the factors that could in theory influence the accuracy of counting molecules by use of quantitative fluorescence measurements are in practice insignificant contributors. The CCD detectors in common use for wide-field microscopy respond linearly to variations in photon input over their entire range up to saturation. The total number of fluorescence photons emitted from an ensemble of FP molecules is, in microscope measurements, proportional to the number of molecules in the assembly. In densely packed assemblies of FP molecules, some degree of “concentration quenching” may occur, reducing the *rate* of fluorescence emission (i.e., apparent brightness), but this does not reduce the cumulative photon yield per molecule before it is photobleached. The primary mechanism for this concentration quenching is thought to be a decrease in the non-radiative lifetime of the excited state, thus reducing the rates of fluorescence emission and photobleaching by the same factor (Hirschfeld, 1976).

One source of non-linearity that can be achieved in some microscopes is ground-state depletion due to the intense excitation coupled with the finite lifetime of the excited singlet or triplet states. However, that is a problem only in point-scanning confocal or 2-photon microscopes. In a wide-field microscope the irradiance is too low to cause significant ground-state depletion. For instance, each mCherry molecule in the experiments reported here is excited and emits a photon approximately once every 10 milliseconds, but its fluorescence lifetime is only 1.5 nanoseconds. Therefore occupancy of the excited singlet state decreases the ground state population by less than one part in 5 million. In most spinning-disk confocal microscopes, the irradiance is also typically too low to cause significant ground-state depletion. However, in single-point scanning confocals, ground-state depletion is easily achieved. Fortunately, it is also easy to detect and easy to avoid (Murray, 2013), so that in principle the fluorescent virus standards could be used in confocal microscopy as well. Multi-photon microscopy is much more problematic, due to the nonlinear and variable relationship between fluorescence emission and photobleaching (Patterson and Piston, 2000).

Milder deviations from the expected strict linear relationship between fluorescence emission and photobleaching are in fact observed for most FPs even with single-photon excitation. The extent of the deviation varies widely, and has been measured for most of the commonly used FPs (Cranfill et al., 2016). The effect is manifested as greater than expected photobleaching when illumination is intense, as in single-point scanning confocals. For this reason, measurements of fluorescence and photobleaching of the virus standards and the cellular target structure need to be carried out using similar illumination intensities. Thus, using wide-field microscopy for the virus standards but confocal microscopy for the cellular target would not give accurate results for many of the FPs (Cranfill et al., 2016).

There are several other sources of inaccuracy that are potentially serious, but can be minimized by careful attention to the measurement and analysis procedures. Significant temporal variations in excitation intensity are nearly universal with mercury arc lamps, and spatial variation across the field of view is also common. Using the proposed “Half-Yield” as the measure of brightness instead of a single time-point measure of fluorescence largely eliminates both of those sources of error. Difficulties in accounting accurately for background fluorescence have a surprisingly large effect on the preciseness of the measurements. Many different computational methods for estimating and subtracting the background fluorescence were explored. The problem is merely a practical one, but nevertheless remains an important limitation on the preciseness of the measurements, whether carried out with a fully automated procedure, or by interactive manual virus identification and measurement. After many trials, the best performance was found to be with a fully automated procedure for the virus images, using the algorithm described in Methods.

The underlying difficulty with any protocol for estimating background is the need to sum the acquired signal over a sufficiently large area to allow for the effect of inescapable small defocus errors. One might think of using only the peak brightness in the image of each virus particle as a measure of its brightness, but at the high NA required for fluorescence imaging of these particles, the peak brightness is exquisitely sensitive to minuscule errors in focus (~20% decrease for 0.1 µm error). The total fluorescence summed over the entire area of the PSF is much less sensitive, but the summation must be taken over a large enough region to include the defocused PSF (at NA 1.3, a defocus error of 0.1 µm approximately doubles the diameter of the PSF). Thus to allow for the possibility (near-certainty) of small random defocus errors, the area of summation must be enlarged by roughly four-fold. Errors in estimating the true background, again a near certainty if the background changes significantly over this spatial scale, are thus greatly magnified in the final result.

The most important sources of inaccuracy when using the fluorescence of FPs to count molecules are unfortunately also the least understood and most difficult to control, namely the complex photophysics of the FP and its sensitivity to local chemical environment (e.g., (Mamontova et al., 2015). Several examples of this were encountered in the work reported here, stemming from the existence of both short and long-lived transient dark states. The good news is that it seems the FP photophysical behavior often suffers from the same eccentricities when incorporated in the virions used for fluorescent standards as when in cellular assemblies. To ensure that these eccentricities are in fact shared, it is important to demonstrate that the “photobleaching” rates for the fluorescent standards and the cellular unknown are the same when measured under identical illumination conditions. If the bleaching rates are closely similar when measured under the same illumination conditions, then one can be reasonably confident that the fluorescent virus particles are an appropriate calibration standard. Note that a large difference in bleach rates does not necessarily mean that the virus particles are inappropriate standards. For instance, a different extinction coefficient for FP in the virus and FP in the cellular organelle would lead to different observed bleach rates, but HY per molecule would be unchanged, so the virus would still provide an accurate calibration for molecular counting. However, since identity in bleach rates provides the only experimental confirmation of the absence of factors that *would* influence HY, one’s confidence in the molecular counts is degraded if the bleach rates for virus and cellular organelle are found to be different. It is common to observe differences in photochemical/photophysical behavior among different FPs used for the same application, as well as among different applications using the same FP (Shaner, 2014). These difference are not yet predictable. Users of the fluorescent virus standards are thus advised to compare several of the seven different FPs available to determine which one performs best in their particular application.

The Sindbis virus fluorescent standards are sub-resolution objects. The two applications described here both involve cellular organelles that are less than 1 μm in diameter, small enough for a good estimate of their fluorescence emission to be obtained from brightness measured in a single “in-focus” image. In a wide-field microscope with high NA objective and commonly used light sources, the total brightness in the image of a fluorescent point object does not change with defocus out to several microns away from the in-focus plane, provided that the summation is done over the entire “geometric shadow”. Thus up to one or two microns in thickness, the size of the cellular organelle can probably be ignored to a first approximation, and images from a single focal plane can be used to measure brightness. However, extension of this method to much larger objects will require confronting some difficult issues in 3D microscopy and 3D deconvolution. In particular, in my experience, the deconvolution algorithms in commonly available software packages perform quite erratically in terms of “photon conservation”: the number of signal photons in the raw images can be substantially different than the number in the final deconvolved result (*data not shown*), thus making quantitative fluorescence measurements on thick objects problematic.

It is necessary to be realistic about the potential accuracy and preciseness of counting molecules in cells using the fluorescence generated from attached FPs. Ideally both preciseness and accuracy would be limited only by the inevitable shot noise in the measured fluorescence, but this is far from the case. As in the examples described here, the statistical uncertainty within one set of measurements can be kept small by careful attention to detail and by averaging over a large number of measurements. A statistical preciseness of the order of 5% for cellular organelles and less than 1% for virus particles is attainable without undue exertion. A somewhat larger variability, of the order of 10%, is observed among nominally identical replicates on different days, using the same hardware but freshly prepared samples each day. By comparison, the accuracy of the measurements can be compromised on a larger scale (almost 2-fold has been observed) if the FP in a cell has different photophysical properties than the FP in the fluorescent standards. At present, the best method for confirming similar photophysical properties is measurement of the bleach rate as described above.

### Uses for the fluorescent virus

Among the motivations for wanting to know the absolute number of molecules of specific proteins, one that has not yet received widespread attention is the use of such numbers as constraints in constructing detailed structural models of large macromolecular assemblies whose overall 3D structure has been determined by electron microscopy. The new direct-electron detectors are revolutionizing structure determination by cryo-electron microscopy, and the size of assemblies whose structure can be determined to sub-nm resolution by this method continues to increase (Zhou, 2011; Grigorieff, 2013; Harapin et al., 2013; Bai et al., 2014). In the past, determining the stoichiometric relationships among the different protein components of macromolecular assemblies often depended on analysis of band intensities in Coomassie-stained SDS polyacrylamide gels. Although those gel-derived stoichiometries were sometimes important in constraining hypotheses about the arrangement of proteins in the 3D structure (e.g., (Amos, 1977)), their use was fraught with difficulties stemming from the uncertain relationship between abundance in cell lysates or subcellular fractions to abundance in specific subcellular organelles and assemblies within cells. Counting molecules based on light microscopic imaging of specific proteins within an intact assembly in a living cell removes those uncertainties, and should thus become a powerful enabling technology for molecular structure determination by cryoEM.

By comparing the fluorescent brightness of conoids in mCherryFP-TgDCX and TgDCX-mNeonGreenFP knock-in lines of *T. gondii* with that of alphavirus particles containing 240 copies of mCherryFP or mNeonGreenFP, I estimated that a conoid contains approximately 2550 TgDCX molecules. The 14 conoid fibers are approximately 430 nm long and contain 9-10 protofilaments, with tubulin dimer spacing of 8 nm along the protofilament, the same spacing as in a canonical microtubule (Hu et al., 2002). Therefore there are 6800 - 7500 tubulin dimers per conoid, or approximately one DCX molecule per 2.6 tubulin dimers. In terms of protein mass, and therefore of scattering power in cryoEM, the contribution from DCX (~ 30 kDa) must be ~10% of that from tubulin (dimer ~ 100kDa), sufficient to give a discernible signal in Fourier transforms from cryoEM images of conoid fibers if the DCX is bound periodically to the tubulin. One possibility is the observed 16 nm periodicity along conoid fibers previously reported (Hu et al., 2002). If DCX is located uniformly along the conoid fibers at 16 nm intervals, then there would not be enough to coat all of the protofilaments. Only approximately 7 of 9, or 8 of 10, protofilaments would be decorated, perhaps leaving bare the protofilaments on the two free edges of the ribbon-like fibers. It will be interesting to compare Fourier transforms from cryoEM images of conoid fibers prepared from these knock-in lines with the transforms of wild-type conoid fibers. The extra mass associated with the FP in the knock-in lines should approximately double the strength of the 16 nm periodicity if this proposed structural arrangement is correct.

Comparing the fluorescence of mCherryFP-Sindbis with fluorescence from TgAPR1-mCherryFP knock-in lines of *T. gondii* led to an estimate of approximately 1250 TgAPR1 molecules in the apical polar ring structure. The 22 cortical microtubules appear to be anchored in the ring, but so far there is no indication of a direct interaction between TgAPR1 and tubulin. At least two other proteins have been observed to localize to the ring (RNG1 (Tran et al., 2010) and RNG2 (Katris et al., 2014)). From genetic evidence, it seems unlikely that TgAPR1 and RNG1 interact (*Murray 2010; Leung 2016; unpublished data*). No information is yet available about interactions of TgAPR1 with RNG2. Unfortunately therefore, appreciation of the significance of the number of molecules of TgAPR1 in the ring must await additional information on its molecular composition.

The applications of the fluorescent virus standards presented in this paper utilized homologous replacement to tag every molecule of a target protein in the cell with an FP. There is no reason the virus particles could not be used as fluorescent standards in other situations, but if the goal is to estimate the total number of molecules of a protein included in a cellular structure, then gene replacement is necessary, as the fraction of molecules tagged with an FP would otherwise not normally be known. On the other hand, in systems where gene replacement is not feasible, counting FP-tagged proteins following transient or stable transfections might provide a useful lower bound on the number of molecules of the target protein in an organelle or other cellular structure.

The fluorescent Sindbis viruses described here have important uses extending beyond their role as calibration standards for counting FP molecules. As described previously (Murray et al., 2007) bright but photobleachable small particles are powerful tools for evaluating performance of microscopes used for 3D fluorescence microscopy. Given the assurance that the particles are all identical, and the ability to generate essentially unlimited quantities on demand, the utility of these tools is enormously increased.

A useful characterization of fluorescent proteins for the purposes of comparing photostability has been proposed (Shaner et al., 2008). In that application, a measurement of the power of the excitation light source is used to give a theoretically predicted photon emission rate per FP molecule, which is then combined with an experimental measurement of “photobleaching” rate (i.e., rate of irreversible fluorophore destruction plus rate of reversible conversion to a long-lived dark state) to calculate a predicted half-time under standardized illumination conditions. This measure is very useful for comparing the photostability of different fluorescent proteins. However, the derived parameter was not designed for use in counting molecules, and it would need to be used with extreme caution in that application, as it is based on a theoretically predicted rather than the actual observed photon emission rate from the FP.

## Acknowledgements

I am grateful to Dr. Richard Day, Indiana University School of Medicine, for providing plasmid DNA encoding several fluorescent proteins, and to Dr. Suchetana Mukhopadhyay for plasmid TE12. I thank Dr. Yu-Chen Hwang and Dr. Kulika Chomvong for assistance in bacterial expression and purification of mCherryFP and EGFP proteins, Dr. Joseph Stampfli, Department of Mathematics, Indiana University, for insights on the aggregate statistical properties of mixtures of different populations, and Drs. Ke Hu, Jacqueline Leung and Jun Liu, Department of Biology, Indiana University, for helpful discussions and critical reading of the manuscript. I am indebted to Dr. Owen Saxton (University of Cambridge) for providing the source code for his “Semper” image processing package. Dr. Jim Powers in the Light Microscopy Imaging Center and Dr. Barry Stein in the Electron Microscopy Center at IU Bloomington provided invaluable assistance with fluorescence microscopy and electron microscopy respectively.

